# Discovery of Waddington’s developmental canals elucidates the embryogenesis stability in *Caenorhabditis elegans*

**DOI:** 10.1101/2023.12.30.573745

**Authors:** Jianguo Wang, Long Xiao, Zeqi Yao, Shanjun Deng, Jianrong Yang, Chung-I Wu, Zhuo Du, Xionglei He

**Affiliations:** State Key Laboratory of Biocontrol, School of Life Sciences, Sun Yat-sen University, Guangzhou 510275, China; State Key Laboratory of Molecular Developmental Biology, Institute of Genetics and Developmental Biology, Chinese Academy of Sciences, Beijing 100101, China

**Author notes:** These authors contribute equally (Jianguo Wang, Long Xiao). Correspondence should be addressed to: X.H. or Z.D.

## Abstract

During embryogenesis, the cells in an embryo need to make numerous spatiotemporal decisions. However, there is inherent noise in each decision due to genetic or environmental fluctuations. How to suppress the noise accumulation to achieve stable embryonic end-products, a process known as Waddington’s developmental canalization, has been a major puzzle in biology since the 1940s. Previous studies have focused on the molecular noise within a cell instead of the cell noise within an embryo, thus providing indirect understandings (e,g., the well-known genetic capacitor Hsp90). In this study, we applied time-lapse microscopic imaging to capturing the spatiotemporal features of single cells, including cell position and cell cycle length, during the embryogenesis of approximately 2,100 *Caenorhabditis elegans* embryos exposed to various genetic or environmental perturbations. By treating the deviation of a cell’s spatiotemporal feature from the expected value as noise, we modeled the transmission of noise from each mother cell to their daughters. We discovered pervasive mother-daughter negative feedbacks, which collectively constitute continuous and comprehensive ‘canals’ for suppressing noise accumulation along the developmental cell lineages, with the steepness (measuring noise suppression efficacy) and depth (measuring noise tolerance level) of the canals quantitatively defined. The learned quantitative rules enabled us to develop a cell-noise-based model that accurately predicts the nematode hatching phenotype, revealing how embryonic stochasticity could cause phenotypic disparity. With a high-dimensional mathematical tool we then proved the system stability of embryogenesis against the cell spatiotemporal noise. We also revealed several dozen canal-maintaining genes and proposed a novel association study framework that links embryonic cells rather than genetic variants with organismal traits. In sum, this study discovered and quantitatively characterized the developmental canals that directly stabilize embryogenesis in a metazoan, illuminating an 80-year-old puzzle and paving a way for studying the phenotypic plasticity and robustness of multicellular organisms from the perspective of embryogenesis.

## Introduction

The developmental programs that decide where and when a cell migrates and divides in an embryo can be highly precise. However, fluctuations arising from genetic, environmental and stochastic factors unavoidably introduce noise, the deviation from the expected value, into each decision. As development is a time series process with an expanding cell population, such noise could quickly accumulate over time and become non-negligible by the end of development, undermining the phenotypic stability of the end-products. Hence, biologists have long been intrigued by the extraordinary stability of developmental outputs observed in most multicellular species^1^.

In his seminal 1942 paper^2^, Waddington wrote, “…*developmental reactions are in general canalized. That is to say, they are adjusted so as to bring about one definite end-result regardless of minor variations in conditions during the course of the reaction*.” Since then, canalization has been a central concept in the fields of developmental biology, evolution, genetics, and systems biology^3-16^. It is often illustrated by a metaphor system in which balls roll through branching canals to their end points; the steeper the canals, the stronger the tendency of the ball to go back to its original course when it is pushed from the course of the canal bottom by internal or external disturbances. Developmental canalization originally referred to the stability of embryonic phenotype^2^, although all levels of molecular networks underlying the embryonic phenotype are expected to be stable as well. Intriguingly, the discovered canalization-associated genes or mechanisms, such as heat shock proteins (e.g., Hsp90)^17^, microRNAs^18^, piRNAs^19^, enhancer redundancy^20^, duplicate genes^21^, genetic compensation^22^, genetic interaction^23^, network buffering^24^, DNA methylation^25^, etc., all function in stabilizing the molecular networks within a cell, with none directly dealing with the cell noise at embryonic level. For instance, the well-known genetic capacitor Hsp90^26,27^, a chaperon protein that suppresses the effects of genetic variants by facilitating the folding of some otherwise misfolded proteins, was first characterized in the yeast *Saccharomyces cerevisiae*, a single-celled organism without embryogenesis^28^. Accordingly, direct understandings^29-33^ on how canalization is achieved at the embryonic level remain limited despite it being 80 years since Waddington coined such an influential concept. In particular, the developmental ‘canals’ for stabilizing embryogenesis have not been quantitatively described in any organism.

Until recently, the lack of ideal data on cell spatiotemporal information during embryogenesis has hindered progress in this area. Recent advances in fluorescence microscopy, single-cell lineage tracing and quantitative phenotyping have made an automated single-cell-resolution quantification of such information possible^32,34-36^. In this study, we used time-lapse microscopy to track the embryogenesis of the nematode *C. elegans* under over 40 environmental conditions^33^. Together with the single-gene RNAi knockdown embryos whose embryogenesis has been tracked by our recent study^33^, a total of ∼2,100 embryos subject to various environmental or genetic perturbations were available. The spatiotemporal features, including cell position and cell cycle length, were recorded for individual cells that appeared in each of the ∼2,100 embryos. Because the development of *C. elegans* follows an invariant cell lineage map to produce a fixed number of cells each with a defined position and function^37^, cells of the same lineage identity in different embryos are expected to have the same spatiotemporal features. Despite substantial between-embryo variations observed for individual cells, the vast majority of the embryos ended up with a phenotypically normal organism, highlighting canalization as an essential component of development. The data we obtained here thus provides a unique opportunity for understanding developmental canalization at the embryonic level.

## Results

### Single-cell-resolution tracking of *C. elegans* embryogenesis using time-lapse microscopy

As in our recent study^33^, we used time-lapse microscopic imaging to track the embryogenesis of *C. elegans* (**Methods**). In brief, the ubiquitously expressed histone::mCHerry fluorescence was used to label individual cells to achieve single-cell-resolution tracking of the nematode embryogenesis from the 4-cell stage up to the 350-cell stage, which corresponds to the completion of nine rounds of cell division (out of ten in total). The spatiotemporal features were recorded for all individual cells that appear within the tracked time window (**Fig. 1a; Methods**). Specifically, the time-lapse three-dimensional positions (*x*, *y* and *z* coordinates) of the cells were available; meanwhile, the cell division timing was available for every cell, with which cell cycle length (CCL) can be readily derived from the time interval between two consecutive cell divisions.

**Fig. 1.**
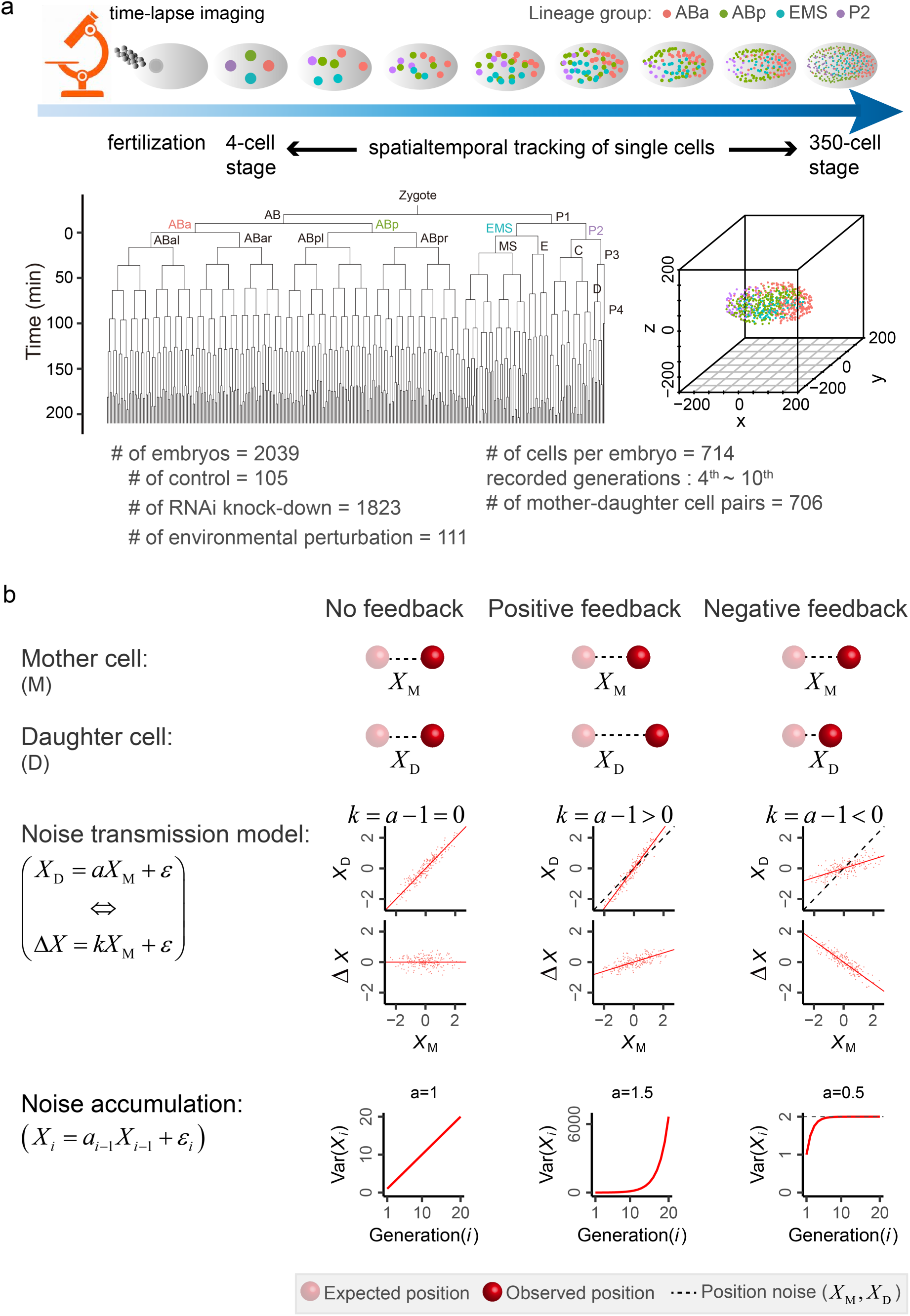
Summary of the Embryogenesis Data and Models for Describing Noise Transmission. **(a)** Time-lapse microscopic imaging is used to track the spatiotemporal features of individual cells, including cell cycle length (CCL) and 3D cell positions (*x*, *y*, and *z* coordinates), from the 4-cell stage to the 350-cell stage during *C. elegans* embryogenesis. A typical *C. elegans* cell lineage tree is presented, with the time axis indicating the cell division timing of each cell, beginning approximately at the 4-cell stage. Accompanying this, the spatial distribution of embryonic cells is displayed, with the four lineage groups (ABa, ABp, EMS, and P2, determined at the 4-cell stage) labeled in different colors. **(b)** Models for describing noise transmission from a mother cell to its daughter. Here, the x-coordinate position noise is considered. When the mother’s noise is fully transmitted to the daughter (i.e., no feedback), a slope of *a* = 1 is expected in the regression analysis between *X*_M_ and *X*_D_ across embryos. This is equivalent to a slope of *k* = 0 between *X*_M_ and Δ*X*, where Δ*X* = *X*_D_ - *X*_M_. Over-transmission (positive feedback) is expected to result in *k* > 0, while incomplete transmission (negative feedback) is expected to result in *k* < 0. Solid and dashed lines represent the regression lines and the diagonal lines, respectively. In addition, we derived the theoretical noise accumulation over generations along a cell lineage. Without loss of generality, we set *a_i_*=1 for all M-D cell pairs of a lineage in the scenario of no feedback, *a_i_*=1.5 in the scenario of positive feedback, and *a_i_*=0.5 in the scenario of negative feedback. The *ε_i_* is set to follow the standard normal distribution. The variance of *X*_i_ (Var(*X_i_*)) is used to assess the noise accumulation over generations. Importantly, Var(*X_i_*) increases with a diminishing return in the scenario of negative feedback, highlighting how negative feedback can prevent noise accumulation.

To mimic environmental perturbations, we exposed nematode embryos to 40 different stress environments, which represent eight types of stressors (temperature, heat shock, bacterial food, stearic acid, glucose, NaCl, FeCl_3_, and paraquat) (**Methods**). In total, we tracked 111 environmentally perturbed embryos (hereafter called Env embryos), all possessing a wild-type genotype, and obtained the aforementioned cell features. To examine the effects of genetic perturbations, we included 1,823 embryos (hereafter called RNAi embryos) that were cultured in a normal environment but each subjected to single-gene RNAi knockdown (with ∼700 different genes involved). The embryogenesis of these RNAi embryos has been tracked by our recent study with the same imaging protocol^33^. In addition, the embryogenesis data of 105 control embryos with neither environmental nor genetic perturbations were also included. Taken together, a total of 2,039 embryos were examined in this study (**Fig. 1a and Table S1-S3**).

Because the developmental cell lineages of *C. elegans* are stereotyped, we can compare the cells of the same lineage identity across different embryos^38^. Considering that some embryos have unusual global sizes or CCLs, we need to normalize the raw cell feature values across different embryos to enhance comparability^29^. To do so, for CCL we linearly aligned each embryo to a randomly selected reference control embryo (ctr-emb58) to derive embryo-specific scaling factors (**Fig. S1; Methods)**; the raw CCL values of an embryo are then all divided by the embryo-specific scaling factor to obtain the scaled CCL values of the embryo. For cell position, we considered the first recorded position of each cell (i.e., the position when the cell was born or birth position). Each embryo was first translated and rotated such that they have matched *x*, *y* and *z* coordinates, with the gravity center of all cells at the point of origin (i.e, gravity center has *x*=0, *y*=0, *z*=0) (**Fig. S2; Methods**). Then, with the same procedure as for scaling CCLs, each embryo was aligned to the reference embryo (ctr-emb58) to obtain the scaled 3D position values (*x*, *y* and *z*) (**Fig. S2; Methods**). The scaling is not sensitive to the choice of reference because the scaling factors obtained based on ctr-emb58 are highly correlated to those based on the average of all control embryos (Pearson’ *R* > 0.999 for CCL, x-coordinate and y-coordinate, and = 0.984 for z-coordinate; **Fig. S1-S2**). For each cell feature of each cell, the mean scaled value is nearly identical for Env, RNAi and control embryos, whereas the standard deviation is substantially larger for Env and RNAi embryos compared to the control embryos (**Fig. S3**). The nearly identical mean values underscore the rationality of the noise definition in the following analyses; meanwhile, the elevated standard deviations among perturbed embryos facilitate the characterization of developmental canalization. Unless otherwise stated, all following analyses in this study are based on the scaled cell feature values.

### Modeling cell noise transmission along developmental lineages

This study examined four cell features, namely the three-dimensional cell positions (*x*, *y* and *z* coordinates) and CCL. Notably, considering the *x*, *y* and *z* coordinates separately would facilitate the definition of both the magnitude and the direction of cell position noise. Using the x-coordinate position as an example, we modeled the position of a daughter cell (*x*_D_) with that of its mother cell (*x*_M_) by an ordinary linear regression function:

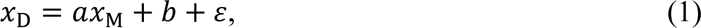

where *a*, *b* and *ε* are slope, intercept and residual error in a linear regression, respectively. This model aligns with previous studies on noise transmission along bacteria or yeast cell lineages^39-41^. Eq. (1) is equivalent to 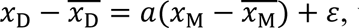 where 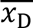 represents the mean x-coordinate position of the daughter among all embryos, 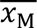 represents that of the mother, and 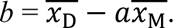 As such, 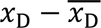 and 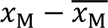 signify the x-coordinate noise of the daughter and the mother, respectively. Hence, as in previous studies^39-41^, the noise we here defined refers to deviation from the mean of the examined population. The noise transmission model can then be written as:

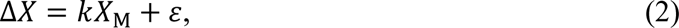

where 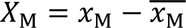 and 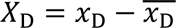 represent the noise of the mother and the daughter, respectively, Δ*X* = *X*_D_ − *X*_M_ represents the noise difference of the daughter from its mother, and *ε* is the same as that in Eq. (1). Notably, the slope *k* in Eq. (2) is equal to *a* -1, where *a* is the slope in Eq. (1).

As shown in **Fig. 1b**, there are three possible scenarios: First, considering that the position of a cell is based on its preceding position, the daughter would fully inherit the position noise of its mother such that *k* = 0 (or *a* = 1) is expected if no feedback is involved. Under this scenario, the noise would accumulate constantly along a cell lineage. Second, there is a positive-feedback regulation wherein the noise of daughter is enhanced relative to that of its mother. Under this scenario, *k* > 0 (or *a* > 1) and the noise would accumulate at a faster rate than the scenario of no feedback. Third, there is a negative-feedback regulation, characterized by *k* < 0 (or *a* < 1), such that only a fraction of the mother’s noise is transmitted to the daughter. Importantly, as what the mathematical principle predicts (**Fig. 1b**), continuous negative feedbacks can effectively prevent noise accumulation along a cell lineage.

### Mother-daughter negative feedbacks for suppressing position noise accumulation

We first looked at ABal-ABala, a randomly selected mother-daughter (M-D) cell pair with ABal being the mother cell and ABala the daughter cell. **Fig. 2a** shows the x-coordinate position noise of ABal (i.e., *X*_ABal_) versus the daughter-mother noise difference (Δ*X=X*_ABala_ – *X*_ABal_) in each of the 2,039 embryos, with *k* = -0.68 observed. The result suggests a negative-feedback regulation on the daughter’s noise by referring to the mother’s noise. In other words, the mother’s noise is transmitted to the daughter but with a mother-dependent offset (i.e., *kX*_M_). Notably, *k* = 0 implies full noise transmission from mother to daughter while *k* = -1 indicates a complete negative feedback with no net noise transmitted from mother to daughter. Here, the observed *k* = -0.68 means a strong but incomplete negative-feedback effect.

**Fig. 2.**
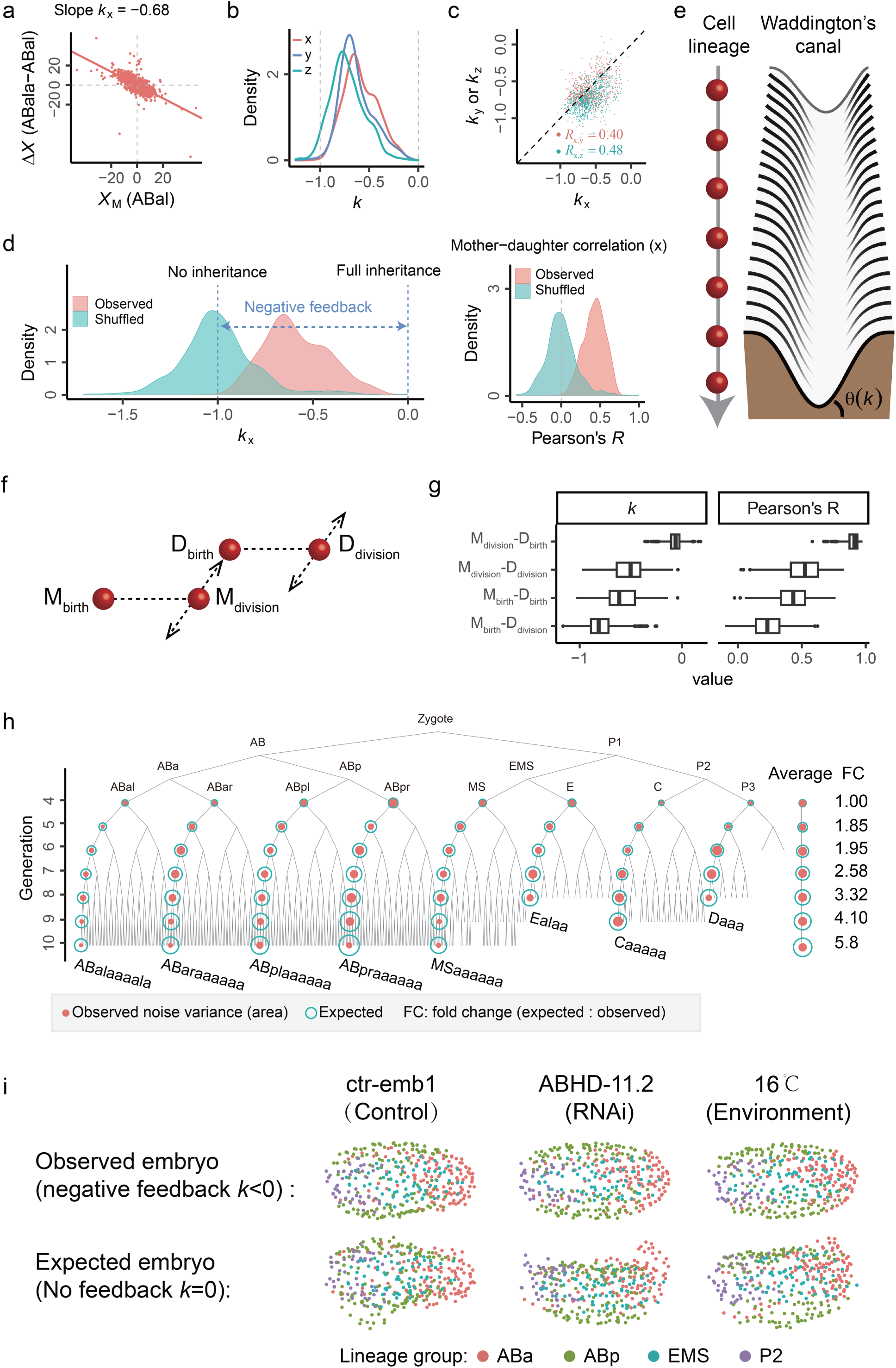
Mother-Daughter Negative Feedbacks for Suppressing Noise Accumulation. **(a)** The scatter plot shows the *X*_M_ and Δ*X* of an example M-D cell pair in 2,039 embryos, with the regression line showing the slope (*k*_x_ = -0.68), which signifies negative feedback. **(b)** The density distribution of *k* values of all M-D cell pairs at *x*, *y*, and *z* coordinates, respectively. The related p-values are available in Fig. S4. **(c)** The *k*_y_ and *k*_z_ are plotted against *k*_x_ across all M-D cell pairs. Pearson’s *R* is used to indicate their correlations. The dashed line shows the diagonal line. **(d)** Pseudo M-D cell pairs are created by shuffling daughters and their mothers, which are then used to derive the expected distribution of *k* and *R* of M-D cell pairs. For shuffled M-D pairs, the *k* values are around -1 and *R* values are around zero, representing no noise inheritance from mother to daughter. Notably, *k* = 0 corresponds to full noise inheritance. For real M-D pairs, the observed *k* values range from -1 to 0, suggesting partial noise inheritance. Consistently, the observed *R* values are larger than zero. **(e)** The ubiquitous negative *k* values indicate continuous negative feedbacks for suppressing noise accumulation along a cell lineage, echoing the concept of Waddington’s developmental canal. In a canal, the ramp always drives a deviating ball back to the bottom of the canal, which resembles negative feedback. Here, the steepness (angle *θ*) of a canal’s ramp, determined by *k* where *θ* = arctan(-*k*), reflects the efficacy of noise suppression. **(f)** In addition to the birth position (the default considered in this study), the division position of each cell can also be analyzed. As such, for each M-D cell pair there are four combinations to be examined. **(g)** The distributions of *k* and *R* values of all M-D cell pairs obtained for the four combinations described in panel **f**. The x-coordinate position noise is considered here. **(h)** Under the null model of noise transmission from mother to daughter (i.e., full noise inheritance), the expected noise variance of a cell equals its specific noise variance (determined by the *ε* in Eq. (1)) plus its mother’s expected noise variance. The expected and observed noise variance are shown for eight representative lineages, labeled in different colors and quantified by the area of the circles. The average variance of the cells of a given generation is also shown, along with the fold change between expected and observed noise variance. **(i)** Three embryos, each corresponding to one of the control, Env, and RNAi embryo classes, are used as examples to show the expected embryonic abnormality by assuming full noise transmission from mother to daughter (i.e., no feedback or *k* = 0). The three embryos developed under real scenarios (i.e., with negative feedback) are shown as comparisons.

To have a general picture we examined all M-D cell pairs at the three coordinates, respectively. The observed *k* is nearly all negative with a strong statistical support (**Fig. 2b, Fig. S4 and Table S4)**, with only four cell pairs being exceptions but just at the y-coordinate. The median *k* is -0.61, -0.66, and -0.74 for the *x*, *y*, and *z* coordinates, respectively. For each M-D cell pair the *k* values at the three coordinates are often similar (**Fig. 2c**). The results suggest nearly ubiquitous negative-feedback regulations on the noise transmission. To simulate a control scenario, we shuffled the cells to create pseudo M-D cell pairs (**Methods**), and failed to find such negative feedbacks (**Fig. 2d and Fig. S5**). Consistently, pseudo M-D pairs in general show no correlation between mother and daughter, indicating no noise transmission from mother to daughter. In contrast, the real M-D pairs often show a weakly positive correlation, a pattern consistent with the negative-feedback model wherein a fraction of mother’s noise is transmitted to the daughter. Interestingly, the ubiquitous mother-daughter negative feedbacks collectively can constitute continuous and comprehensive ‘canals’ for suppressing the position noise accumulation along the developmental cell lineages in *C. elegans*, where the *k* serves as a quantitative measure of the noise suppression efficacy or ‘steepness’ of the canals (**Fig. 2e**). As such, Waddington’s developmental canals are no longer a metaphor but a mathematically formulated reality in the nematode.

Several technical factors could potentially confound the estimation of *k*. First, biases might be introduced by the scaling process. We re-estimated *k* for embryos with different scaling factors and obtained essentially the same results (**Fig. S6**). Second, embryos with a low alignment quality could introduce errors. We focused on the embryos with a high alignment quality and used them to re-estimate *k*; the re-estimated *k* is highly similar to the original *k* (**Fig. S7; Methods**). Third, the ‘dilution effect’ (or the effect of regression towards the mean) arising from measuring error could bias the estimation of *k*. Here, the measuring error of cell position stems from the time interval between two consecutive images taken by the time-lapse microscopy. We estimated the variance caused by the measuring resolution, recalculated *k* after adjusting for the dilution effect, and found the results qualitatively unchanged (**Fig. S8**). Fourth, lineage tracing errors, estimated to be less than 1% in our previous study^33^, might also introduce biases. We simulated lineage tracking errors in the embryos at a rate of up to 10% in each generation and found the estimation of *k* robust **(Fig. S9; Methods**). Fifth, we found the standard deviation of the noise to be comparable between the mother and the daughter of an M-D pair (**Fig. S10**), suggesting that the observation of *k* < 0 in nearly all M-D pairs is not simply explained by a smaller noise variance for daughters than their mothers. Lastly, by generating simulated embryos we confirmed that the ‘translation-rotation-scaling’ alignment procedure had negligible effects on the estimation of *k* (**Fig. S11; Methods**).

To elaborate the process of noise transmission, we may analyze both the birth position (the default position considered in this study) and the division position (i.e., the position when a cell divides) of each cell (**Fig. 2f**). As such, for each M-D cell pair there are four possible combinations to be analyzed, namely, M_birth_-D_birth_, M_birth_-D_division_, M_division_-D_birth_, and M_division_-D_division_, where M_birth_ and M_division_ stand for the birth and division positions of the mother, and D_birth_ and D_division_ the birth and division positions of the daughter, respectively. As shown in **Fig. 2g** and **Fig. S12**, the obtained *k_x_* values tend to be smaller in the combinations with a larger M-to-D time interval (**Table S4**). For instance, the median *k_x_* was -0.06 in M_division_-D_birth_, a combination with a very short M-to-D time interval; meanwhile, the number was -0.81 in M_birth_-D_division_, a combination with an M-to-D time interval spanning nearly two cell generations. Consistently, the correlation between mother and daughter was the largest (median Pearson’s *R_x_* = 0.91) in M_division_-D_birth_ and smallest in M_birth_-D_division_ (median Pearson’s *R_x_* = 0.23). These results convey three notable messages: First, there is nearly full noise transmission observed (in M_division_-D_birth_), which can serve as another control for the signals in **Fig. 2b**. Second, there is minimal net noise transmitted between stages of nearly two cell generations apart (from M_birth_ to D_division_), highlighting the efficacy of negative feedbacks in suppressing noise accumulation. Third, the reduction of net noise transmission does not just occur at the transition moment between mother division and daughter birth; instead, it likely occurs continuously as a function of the time between two examined stages. Accordingly, the signals in **Fig. 2b** represent the cumulative negative-feedback effects over a whole cell generation (from M_birth_ to D_birth_). Of note, the observed reduction of net noise transmission over time cannot be explained by the growing random noise accumulated in the daughter over time. This is because, as what the Eq. (1) predicts, random noise in the daughter would only affect the Pearson’s *R* but not *k* (**Fig. S13**).

What if such negative feedbacks do not exist? In that scenario, *k* = 0 and the noise of a daughter cell would comprise two parts: one inherited fully from its mother and the other representing the daughter-specific noise arising independently from its mother. As a consequence, cell noise would accumulate constantly along a cell lineage by adding up the initial mother cell noise and all subsequent daughter-specific noises. Under the assumption that the *ε* in Eq. (2) can accurately represent daughter-specific noise, we estimated the theoretical cell noise accumulation without negative feedbacks (i.e., *k* = 0) (**Methods**). Due to incomplete data for earlier cells, we considered the eight cells of the 4^th^ generation as the initial mother cells of the lineages. **Fig. 2h** shows the observed and expected (or theoretical) noise accumulations along eight representative lineages. On average, the expected noise variance among embryos is 5.8, 6.6, and 7.7 times the observed noise variance for cells of the 10^th^ generation at the *x*, *y*, and *z* coordinates, respectively (**Fig. 2h and Fig. S14**)). Accordingly, under the scenario of no negative feedbacks the cells would gradually deviate from their pre-determined positions, resulting in morphologically abnormal embryos as shown in **Fig. 2i** (**Methods**). These being said, we noted that this part relied heavily on the assumption that the *ε* in Eq. (2) accurately represents daughter-specific noise.

### Altered negative feedbacks by specific perturbations

In the preceding analyses, embryos of different classes were collectively used to estimate *k*. Next, we considered the Env embryos, RNAi embryos and control embryos separately, re-estimating *k* for each of the three classes of embryos, respectively (**Methods**). As shown in **Fig. 3a-b** and **Fig. S15**, *k*_control_ (i.e, the *k* obtained by considering only the control embryos) is highly correlated with *k*_Env_ (Pearson’s *R* = 0.64, *p* < 2.2×10^-16^) and *k*_RNAi_ (Pearson’s *R* = 0.80, *p* < 2.2×10^-16^), suggesting shared negative-feedback regulations in response to the distinct noise-producing factors. Interestingly, by plotting the *k*_control_-*k*_Eny_ versus *k*_control_-*k*_RNAi_, we found over one hundred M-D cell pairs with a similarly altered *k* in Env and RNAi embryos compared to the counterpart *k*_control_ (**Fig. 3c and Fig. S15; Methods**). Most of these cell pairs (92%, 84% and 77% for *x*, *y* and *z* coordinates, respectively) exhibited an increased *k*, indicating an attenuated negative-feedback effect. The affected cells show an enrichment in two tissues, pharynx and intestine (**Fig. 3d**; **Methods**). Hence, genetic and environmental factors appeared to modulate embryogenesis by altering the noise transmission in specific cell lineages.

**Fig. 3.**
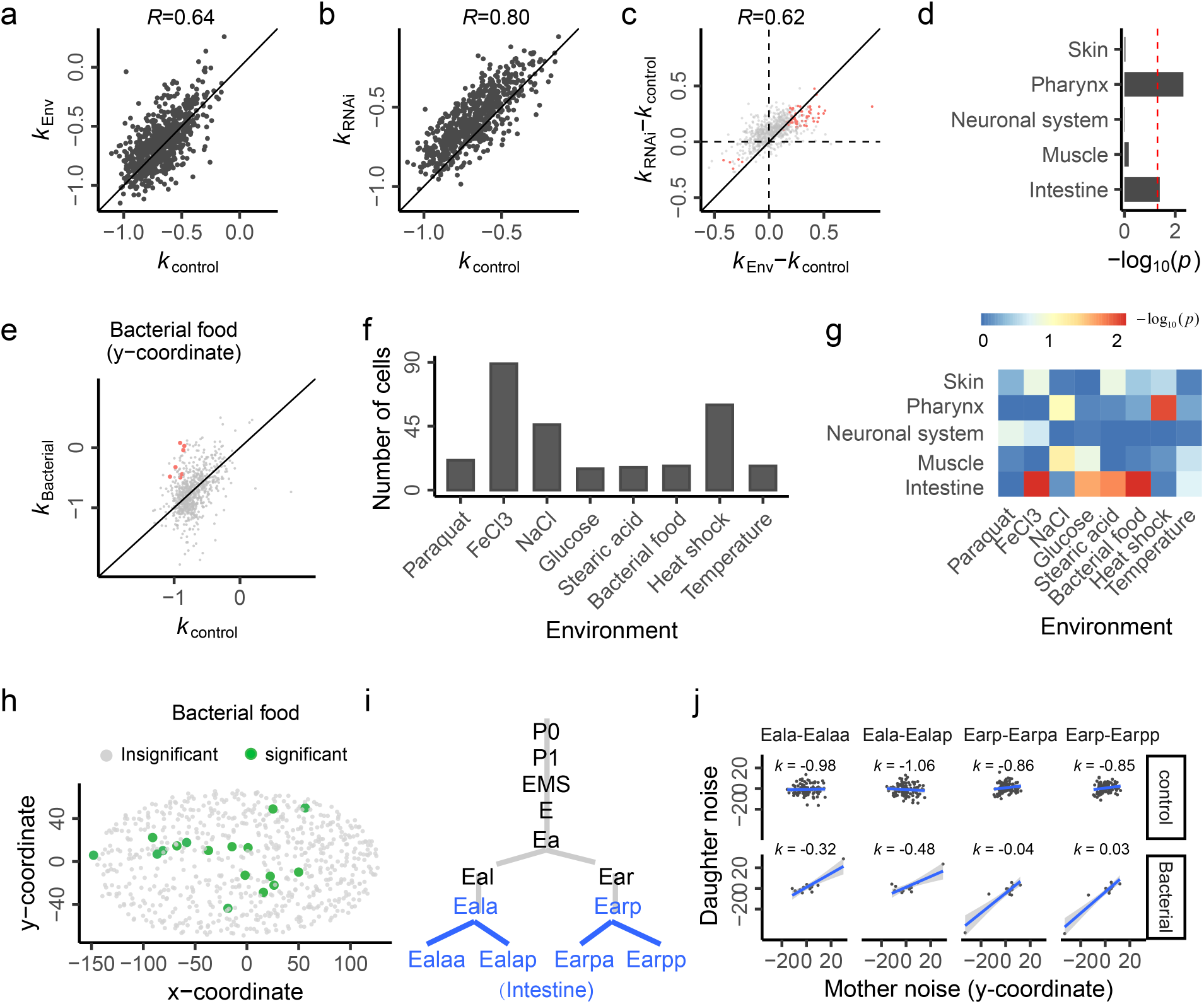
Altered Negative Feedbacks by Specific Perturbations. **(a)** The *k* values of all M-D cell pairs estimated in Env embryos are correlated with those in control embryos (Pearson’s *R* = 0.64, *p* < 2.2×10^-16^). Each dot represents a cell pair and the x-coordinate position noise is considered. **(b)** The *k* values of all M-D cell pairs estimated in RNAi embryos are correlated with those in control embryos (Pearson’s *R* = 0.80, *p* < 2.2×10^-16^). Each dot represents a cell pair and the x-coordinate position noise is considered. **(c)** The difference between *k*_Env_ and *k*_control_ is plotted against that between *k*_RNAi_ and *k*_control_, with Pearson’s *R* shown. The M-D cell pairs with a significant (*p* < 0.05) difference both between *k*_Env_ and *k*_control_ and between *k*_RNAi_ and *k*_control_ are highlighted in red. The x-coordinate position noise is considered. **(d)** Tissue enrichment of the cells with an altered *k* in perturbed embryos, with the dashed line showing *p* = 0.05. After combining the data of three coordinates, there are 125 affected cell pairs with consistently altered *k* in both Env and RNAi embryos compared to the counterpart *k*_control_. For each M-D cell pair the daughter’s cell fate is considered. **(e)** There are seven cell pairs with a significantly (*p*_adjust_ < 0.05) altered *k* at the y-coordinate in embryos exposed to stressful bacteria food. **(f)** The number of significantly affected cell pairs for each environmental stressor type. **(g)** Tissue enrichment pattern of the significantly affected cell pairs of each stressor type. **(h)** The embryonic distribution in the x-y plane of the 17 significantly affected cell pairs (only daughters are shown) under stressful bacteria food. The pairwise Euclidean distance in three-dimensional space within the affected cells (D_within_) is significantly smaller than the distance between the affected cells and other cells (D_between_), suggesting a non-random pattern (one-tailed Wilcoxon test, *p* = 3.52×10^-4^). **(i)** The lineage information of the four cell pairs with a tissue destination of intestine and affected by stressful bacteria food in the noise transmission at y-coordinate. **(j)** The y-coordinate noise in the control embryos versus the bacterial-food-perturbed embryos for the four cell pairs highlighted in panel **i**.

To further demonstrate the phenomenon, we examined specific environmental stressors. The 111 Env embryos are exposed to 40 environments of eight types of stressors, including temperature, heat shock, bacterial food, stearic acid, glucose, NaCl, FeCl_3_ and paraquat, with ∼15 embryos available for each stressor type. We estimated *k* for the embryos of each stressor type, and identified the M-D cell pairs with a significantly (*p*_adjust_ < 0.05, Benjamini-Hochberg adjustment) altered *k* compared to the counterpart *k*_control_ (**Table S5; Methods**). **Fig. 3e** shows the *k* estimated for the 14 embryos exposed to stressful bacterial food, with seven significantly affected cell pairs revealed at the y-coordinate. The total number of significantly affected cell pairs varies substantially across the different stressor types, ranging from 15 (glucose) to 89 (FeCl_3_) (**Fig. 3f**). As shown in **Fig. 3g**, the affected cells in different stressor types show different tissue enrichment patterns, although intestine appears to be generally sensitive to the environmental stressors. Using the 17 significantly affected cell pairs in stressful bacteria food as a showcase, we plotted the embryonic positions of these cells and found them distributed in a non-random pattern (*p* = 3.52× 10^-4^, one-tailed Wilcoxon test; **Fig. 3h**). Among the 17 cell pairs four have a tissue fate of intestine, and the related developmental lineages are shown in **Fig. 3i**. For each of the four cell pairs we plotted their noises in the control embryos versus the bacteria-food-perturbed embryos, respectively (**Fig. 3j)**. There is little mother-daughter correlation observed, with an estimated *k* close to -1, in the control embryos; meanwhile, often a strong positive correlation is observed, with a much larger *k*, in the bacterial-food-perturbed embryos. Hence, the noise suppression via negative feedbacks can be precisely terminated at specific cell lineages by specific environmental factors, which suggests an interesting angle for studying the development plasticity under various environments.

### Early cell noises predict hatching phenotype

The process of embryogenesis in *C. elegans* begins with a zygote and concludes with an organism composed of 558 cells^37^, which then hatches. For the 2,039 embryos examined in this study, we recorded the cell spatiotemporal features from the 4-cell to 350-cell stages as well as the hatching phenotype, namely, hatched (n = 1930) or lethal (n = 109) (**Fig. 4a and Table S6**). We hypothesized that the cell position noises in an embryo could predict the hatching phenotype. We trained a cell-noise-based logistic model within a machine-learning framework and found it can well predict the hatching phenotype, with an area under the curve (AUC) of 0.91 (**Fig. 4b; Methods**). To simulate a control, we shuffled the hatching phenotypes among embryos and obtained an AUC of approximately 0.5, suggesting the good model performance is not explained by hatched-lethal sample imbalance or other technical biases (**Fig. 4b; Methods**). The 53 cells (55 cell features) with a non-zero coefficient in the model, designated as effect cells, were found in diverse lineages at both early and late generations (**Fig. 4c**). To test the prediction performance of early cells, we restricted the predictors by excluding cells appeared after a specified cell generation. We observed rather good prediction performance by using only early cells as predictors (**Fig. 4d**). For instance, the AUC reached 0.71 by considering only the 24 cells of the 4^th^ ∼5^th^ generations, and 0.78 by considering only the 56 cells of the 4^th^ ∼ 6^th^ generations (**Fig. 4d**). The results offer a quantitative understanding on the extent to which the early embryonic noises determine a late organismal phenotype.

**Fig. 4.**
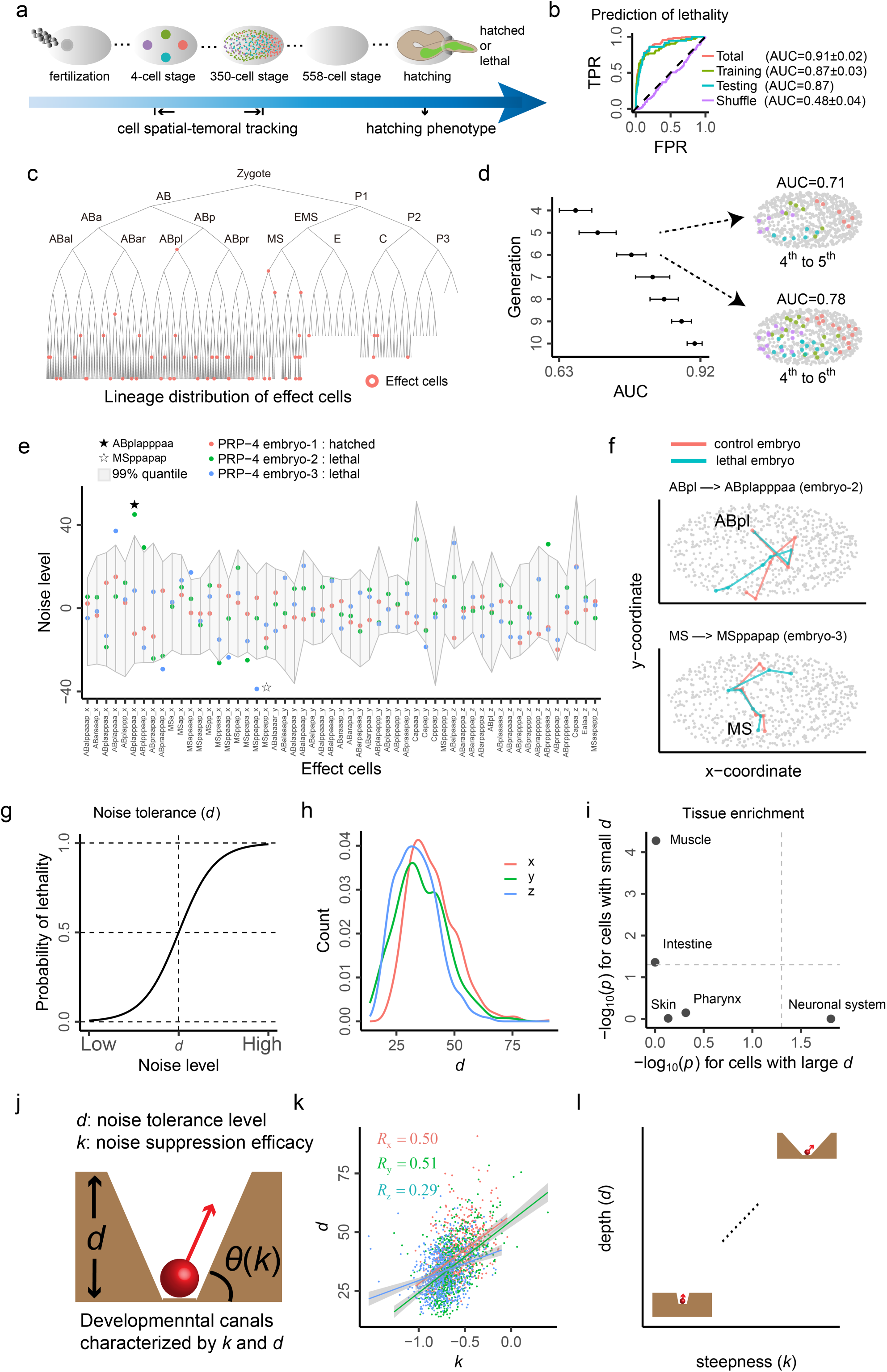
Noise-Based Prediction of Hatching Lethality Characterizes Developmental Canals. **(a)** Following the lineage tracing from the 4-cell to the 350-cell stage, the hatching phenotype of each embryo is recorded, designated as either hatched or lethal. **(b)** By implementing logistic regression under the machine-learning framework of LASSO, the hatching phenotype is modelled by the cell noises. All the embryos are divided into a training and testing set, accounting for 1/3 and 2/3, respectively, ensuring that different embryos of the same RNAi gene are categorized into the same set (either training or testing set, but not both). The AUC is used as a measure of prediction performance. In addition, a logistic model is implemented based on all embryos to obtain the final model for defining effect cells. To evaluate the potential effect of an imbalanced hatched-lethal ratio, a logistic model is also conducted by shuffling the hatching phenotype. The ROC curves along with the AUC (±standard error) are shown for the training, testing, and total sets, as well as the shuffled data. **(c)** The distribution of effect cells (cells with non-zero coefficients in the logistic model) in the lineage tree. **(d)** A series of logistic learning models are implemented by excluding cells after a given generation. The AUCs with standard error are plotted for each generation, demonstrating the predictability of early-generation cells on hatching phenotype. The embryonic distribution of early-generation cells is shown for two example generations. **(e)** For the three embryos subject to the same gene (PRP-4) RNAi, one hatches normally while the other two have a lethal hatching phenotype. Through comparison, the effect cells’ features of the hatched embryo have an overall lower noise level. The grey polygonal region covers the 99% quantile of cell noise among all embryos, within which all the cell features’ noise levels of the hatched embryo are located. Two outlier noise levels are highlighted by five-pointed stars for each of the two lethal embryos. **(f)** The lineage trajectories leading to the two cells highlighted in panel **e** are depicted in the two lethal embryos, respectively. The initial cells are ABpl and MS, respectively. The lineage trajectory of the control embryos and the background cells are derived from the average of the control embryos. For simplicity, only the x-y plane is shown. **(g)** The definition of noise tolerance level (*d*) of each cell. The model between a cell feature’s absolute noise (|*X|*) and hatching phenotype is modeled by logistic regression logit(P) = E(*β*_0_+*β*_1_|*X|*), where P represents the probability of hatching lethality. As such, *d* = -*β*_0_/*β*_1_, corresponding to 50% probability of hatching lethality. **(h)** The distribution of *d* of all cells at each of the three coordinates, respectively. **(i)** Tissue enrichments of the cells with very large or small (top or bottom 5%) *d* values. The *p*-values are calculated by a binomial test. **(j)** An illustration of Waddington’s developmental canals characterized by two parameters *k* and *d*. The *θ*(*k*) is same as that in Fig. 2e. **(k)** There is an overall positive correlation between *k* and *d* (Pearson’s *R* = 0.50, 0.51, and 0.29, for *x*, *y* and *z* coordinates, respectively, with the corresponding *p* < 2.2×10^-16^, *p* < 2.2×10^-16^, and *p* = 7.30×10^-15^, respectively). Each dot represents a cell, and the *k* estimated for an M-D cell pair is assigned to the daughter cell. **(l)** An illustration of the characteristics of Waddington’s developmental canals, with weak negative feedback accompanied by high noise tolerance level (top-right) and strong negative feedback accompanied by low noise tolerance level (bottom-left).

The good performance of the cell-noise-based model sheds lights on an interesting observation that individuals subject to the same perturbation express distinct hatching phenotypes. For example, among the three PRP-4 RNAi embryos, one hatched a viable larva but the other two did not. In line with the divergence of hatching phenotype, the noise levels of the effect cells in the hatched embryo are overall lower than those in the two lethal embryos (**Fig. 4e**). In the hatched embryo (embryo-1), all effect cells have a noise level within the 99% quantile of the 2,039 embryos, while the two other lethal embryos each has a couple of outlier cells with an exceptional level of noise (**Fig. 4e**). A close examination of the lineage leading to ABplapppaa, an outlier cell in embryo-2, and the lineage leading to MSppapap, an outlier cell in embryo-3, revealed the gradual accumulation of noise to become exceptionally deviated from the average cell position of the control embryos (**Fig. 4f**). This highlights the importance of suppressing noise accumulation to achieve successful developmental end-products. Notably, differences between the two lethal embryos are also substantial. For instance, the cell ABplapppaa has an exceptional level of noise in embryo-2 but very small noise in embryo-3 (**Fig. 4e**). The results demonstrate vividly the connection between embryonic stochasticity and phenotypic heterogeneity.

### Charactering the noise tolerance level or ‘depth’ of developmental canals

Since the hatching lethality can be explained by cell noises, we could derive the threshold of noise tolerance for each of the cells. For each cell we defined the noise tolerance level, designated as *d*, as the absolute noise level corresponding to a 50% probability of hatching lethality (**Fig. 4g, Fig. S16 and Table S7; Methods**). Although the length of A-P axis (x-coordinate) is approximately twice that of the L-R axis (y-axis) and three times that of the D-V axis (z-coordinate), the obtained *d* of the x-coordinate is comparable to that of the other two coordinates (**Fig. 4h**). This suggests the absolute noise level, rather than the relative noise level, matters. We checked the cells with a very large (top 5% for any coordinate) or small (bottom 5% for any coordinate) *d* to check their tissue enrichment, respectively (**Methods**). Those with a small *d* are enriched in muscle, suggesting the exact positions of muscle cells are critical for hatching (**Fig. 4i**). Meanwhile, those with a large *d* are enriched in neural system, suggesting a relaxed requirement for the positions of neural cells.

The availability of *d* suggests another parameter to describe the developmental canals that suppress noise accumulation. Specifically, *d* can serve as a measure of the noise tolerance level or ‘depth’ of the canals (**Fig. 4j**). Interestingly, *d* is positively correlated with *k*, the noise suppression efficacy or steepness of the developmental canals (Pearson’s *R* = 0.50, 0.51, and 0.29, for *x*, *y* and *z* coordinates, respectively, with the corresponding *p* < 2.2×10^-16^, *p* < 2.2×10^-16^, and *p* = 7.30×10^-15^, respectively; **Fig. 4k**). This indicates that cells receiving a lot noise from their mothers (i.e., with a large *k*) tend to have a high noise tolerance level, which seems to be a sensible design for the system (**Fig. 4l**). Hence, it is likely that the developmental canals have been shaped by natural selection, instead of being an inherent product/property of a complex system^9^.

We further examined the fluctuation of *k* and *d*, respectively, across cell generations. Overall, the noise tolerance level *d* tends to be larger at earlier generations, suggesting more room for noise at the beginning of development (**Fig. S17a**); meanwhile, *k* appears to be the largest at the middle generations, indicating stronger negative feedbacks at both the beginning and the end of development (**Fig. S17b**). We also looked at the spatial distribution of *k* and *d*, respectively, and observed various non-random patterns (**Fig. S17c-d**). Further studies are required for understanding how such patterns are related to the formation of body structure of *C. elegans*.

### Assessing the system stability of embryogenesis against position noises

The above analyses considered only the marginal effects of mothers on noise transmission, which is not sufficient for assessing the stability of embryogenesis as a dynamic cell system. It is desirable to have a high-dimensional model for describing the noise transmission in the system. We thus expanded Eq. (2) to be

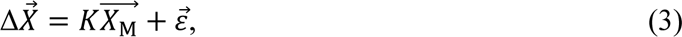

where 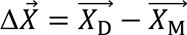 is an *n* × l column vector storing the x-coordinate noise difference of all mother-daughter cell pairs (here *n* = 706), 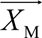 is an *n* x l column vector storing the x-coordinate noise of mothers, 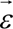 is an *n* x l column vector storing the residual errors, and *K* is an *n* × *n* coefficient matrix with element *K* (*i*, *j*) characterizing the effect of the *j*^th^ mother on the noise difference of the *i*^th^ mother-daughter pair (**Fig. 5a; Methods**). The high-dimensional noise transmission models of the *y* and *z* coordinates were derived in the same way.

**Fig. 5.**
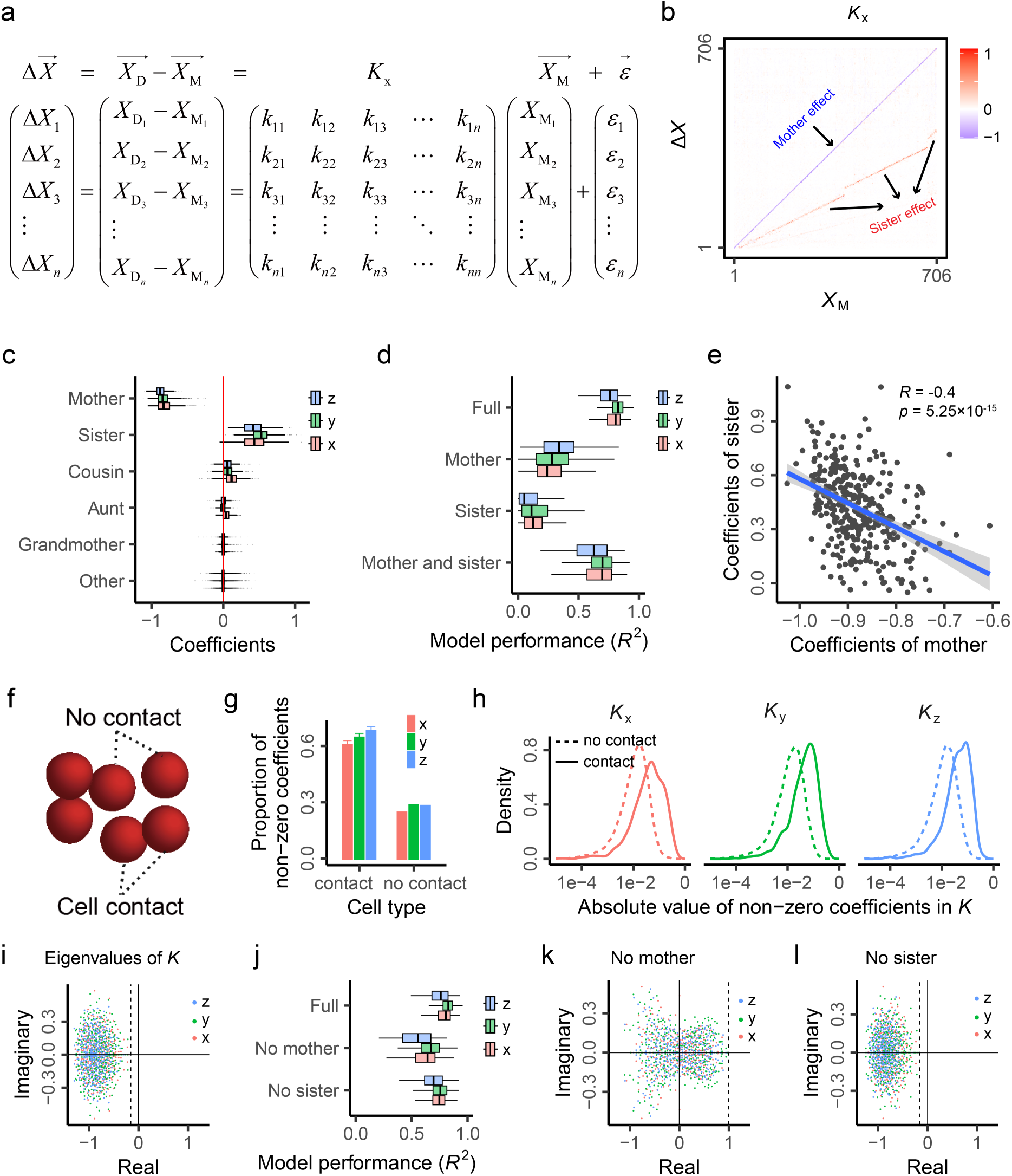
Assessing the System Stability against Cell Position Noises. **(a)** Details of the high-dimensional noise transmission model described in Eq. (3). **(b)** The obtained coefficient matrix *K*_x_ shown as a heatmap. The diagonal coefficients show the mother effects. The mother and sister effects are highlighted in different colors. **(c)** Summary of the coefficients in *K*_x_, *K*_y_ and *K*_z_, with standard boxplots shown. **(d)** The predicting performance of the full model and three nested models that consider only mother, only sister, or both mother and sister, respectively. The model performance is measured as the explained variance (*R*^2^). A standard boxplot is used to show the *R*^2^ of each item in 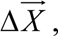 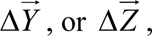 respectively. **(e)** There is overall a negative correlation between mother effect and sister effect in *K*_x_ (Pearson’s *R* = -0.40, *p* = 5.25×10^-15^). The fitted line shows the 95% confidence bands. **(f)** An ideograph showing cells with or with no contact. **(g)** The noise difference of a focal cell from its mother (Δ*X*, Δ*Y*, or Δ*Z*) is more affected by cells with contact to the focal cell, evidenced by the higher proportions of non-zero coefficients of these cells in *K*_x_, *K*_y_ and *K*_z_, respectively. Error bar shows two standard errors. The mother and sister of each focal cell are excluded from the analysis. **(h)** Consistent with panel **g**, the absolute values of non-zero coefficients in *K*_x_, *K*_y_ and *K*_z_ are overall larger for the cells with contact to the focal cell than those of no contact. The mother and sister of each focal cell are excluded from the analysis. **(i)** The eigenvalues of *K*_x_, *K*_y_ and *K*_z_ all have a negative real part, indicating the inherent stability of the dynamic cell system. The x-axis and y-axis represent the real and imaginary parts, respectively, of an eigenvalue. The dashed line marks the maximum real part of the eigenvalues. **(j)** The predicting performance of the full model and two nested models that block either mother or sister. **(k)** The eigenvalues of *K*_x_, *K*_y_ and *K*_z_ in the nested models blocking mother. Many eigenvalues have a positive real part, suggesting the loss of stability after blocking mother. **(l)** The eigenvalues of *K*_x_, *K*_y_ and *K*_z_ in the nested models blocking sister. All eigenvalues have a negative real part, suggesting the system stability remains after blocking sister.

We estimated *K* using linear regression with regularization (**Methods**). Importantly, to model the noise difference of a mother-daughter pair, the focal daughter cell, if present in 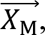 will be blocked by setting the corresponding element in *K* to be zero (**Methods**). When the focal cell is not influenced by other cells except its mother, the high-dimensional model will be degenerated into Eq. (2). As shown in **Fig. 5b** and **Fig. S18a-b**, the coefficient matrix *K* for the three coordinates *(K*_x_, *K*_y_ and *K*_z_) is all sparse. The noise difference of a focal cell from its mother is explained mostly by its mother and sister (**Fig. 5c-d)**, with a negative correlation of their coefficients observed (Pearson’s *R* = -0.40, -0.20 and -0.20 for *x*, *y* and *z* coordinates, respectively, with the corresponding *p* = 5.25×10^-15^, *p* = 1.59×10^-4^ and *p* = 1.99×10^-4^, respectively; **Fig. 5e and Fig. S18c-d**). Some cells within an embryo have intimate cell-cell contacts^42^ with each other (**Fig. 5f**), which may affect their position noise. Indeed, we found that the cells having intimate contact with a focal cell tend to show a stronger effect on the focal cell (**Fig. 5g-h; Methods**).

In linear transformation (here represented by *K*_x_*, K*_y_ or *K*_z_), an eigenvector points to a direction in which it is stretched by a factor of the corresponding eigenvalue. When the real part of an eigenvalue is negative, the direction of the transformation will also be negative, which is analogous to the negative *k* value in Eq. (2). When all eigenvalues of a linear transformation have a negative real part, the examined system can be regarded as stable. Notably, eigenvalue analysis has been widely used to assess the stability of ecological systems^43,44^ and was recently applied by us to studying gene regulatory networks^45^.

We calculated the eigenvalues of *K*_x_, *K*_y_ and *K*_z_, respectively. Remarkably, they all have a negative real part (**Fig. 5i**). This suggests that the position noises are counteracted during the along-lineage transmission in all directions of the cell system. In other words, the dynamic cell system appears to be inherently stable against the position noises. We further tested two models, one with mothers blocked and the other with sisters blocked (**Methods**). While blocking sisters primarily maintain the model performance and system stability, blocking mothers substantially reduces the model performance and undermines system stability (**Fig. 5j-l**). To exclude the potential effects from descendants, we also blocked for each mother-daughter pair the cells of the descendant generations and found largely the same results (**Fig. S19; Methods**). In addition, to mimic the classical model of assessing the stability of a complex system, we examined the noise differences of the same cells between cell birth and cell division and obtained consistent system stability (**Fig. S20; Methods**). These results notwithstanding, we cautioned that there are substantial variances unexplained by the high-dimensional models (**Fig. 5d**), which may confound our assessment of the system stability.

### Suppression of CCL noise accumulation

We next analyzed the noise of cell cycle length (CCL) using largely the same procedure as for analyzing cell position noise (**Methods**). For all examined mother-daughter cell pairs, the slope *k* is invariably negative, ranging from -1.13 to -0.12 with a median of -0.85 (**Fig. 6a, Fig. S21 and Table S4**). This is consistent with a previous observation of small but significant positive correlations between mother and daughter in CCL^29^. The estimated *k* is not biased by the confounding factors considered in the preceding analyses on position noise (**Fig. S21**). Notably, unlike cell position with an apparent mother-daughter inheritance, CCL may or may not have such inheritance. Hence, there may be a molecular ‘modifier’ of CCL that is transmissible from mother to daughter such that the mother’s noise is transmitted to the daughter with a mother-dependent offset (i.e., negative feedback). Alternatively, mother and daughter might each acquire their CCL noise separately despite a shared noise component involved due to unknown reasons; as such, no negative feedback exists. We found the obtained *k* of CCL noise is correlated with the *k* of position noise (Pearson’s *R* = 0.14, 0.39 and 0.23 for *x*, *y* and *z* coordinates, respectively, with the corresponding *p* = 7.25×10^-3^, *p* = 2.83×10^-14^ and *p* = 1.37×10^-5^, respectively; **Fig. 6b**). This observation aligns with the first interpretation, suggesting shared negative-feedback regulations on the CCL noise and position noise. Regardless of the mechanistic interpretations, observation of ubiquitously negative *k* indicates a comprehensive suppression of the CCL noise accumulation. We also calculated the CCL noise tolerance level (*d*) for each of the cells (**Fig. 6c, Fig. S22 and Table S7; Methods**) and observed similarly a positive correlation between *k* and *d* (Pearson’s *R* = 0.30, *p* = 1.44×10^-8^; **Fig. 6d**). In addition, we found integrating CCL noise with position noise improved the cell-noise-based prediction of hatching phenotype, increasing the AUC from 0.91 to 0.96 (**Fig. 6e-f; Methods**).

**Fig. 6.**
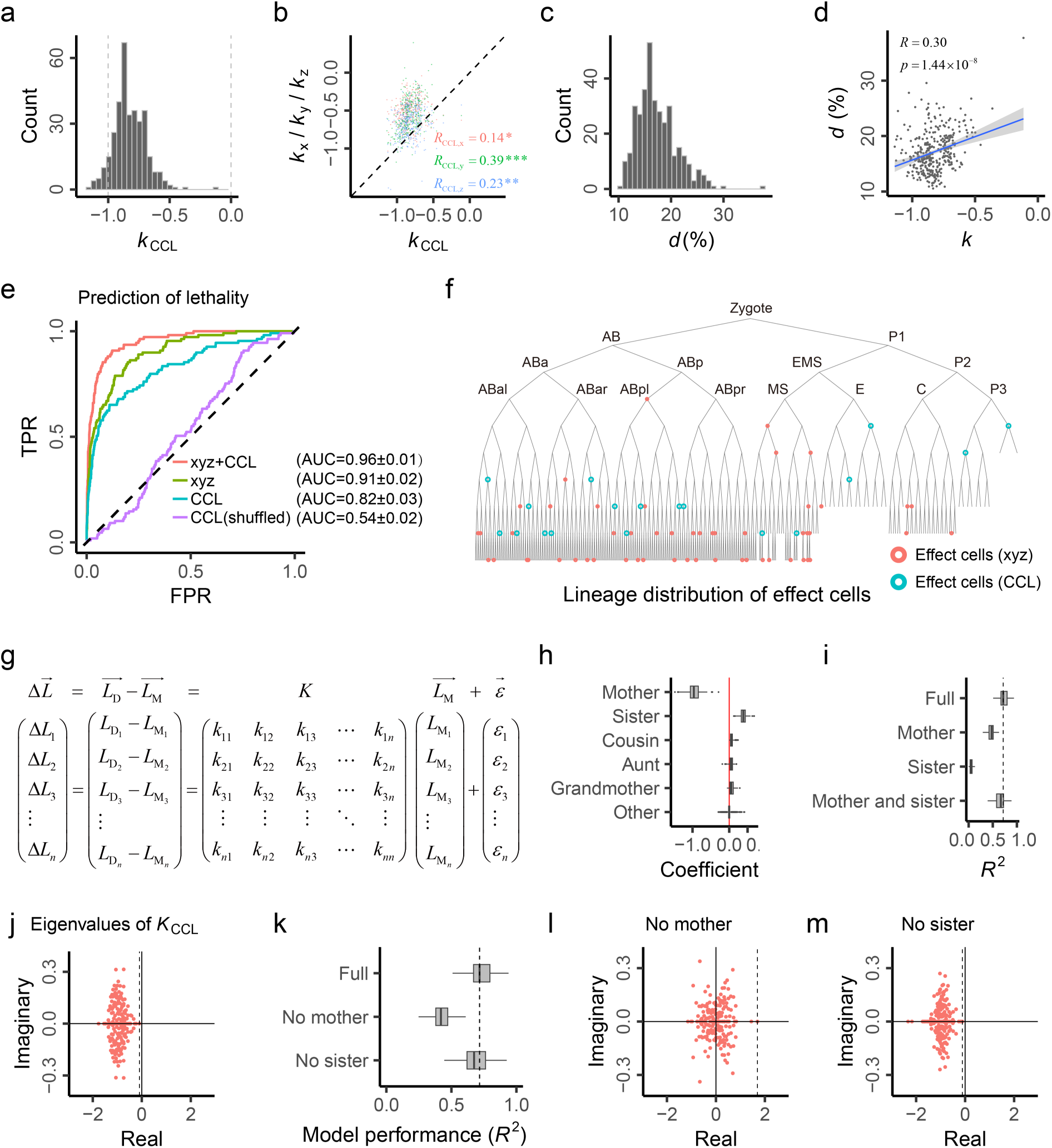
Noise Suppression in Cell Cycle Length. **(a)** The *k* in cell cycle length (CCL) is negative for all M-D cell pairs. The dashed lines show *k* = -1 and *k* = 0, respectively. **(b)** The *k* of CCL (*k*_CCL_) is significantly correlated to *k*_x_, *k*_y_ and *k*_z_, respectively. Pearson’s *R* is shown (*: *p* = 7.25×10^-3^; **: *p* = 1.37×10^-5^; ***: *p* = 2.83×10^-14^). Each point represents an M-D cell pair. **(c)** The distribution of CCL noise tolerance level (*d*) of all cells. Notably, here CCL noise measures the deviation from the average of all embryos divided by the average. **(d)** There is an overall positive correlation between *k* and *d* in CCL (Pearson’s *R* = 0.30, *p* = 1.44 ×10^-8)^. Each dot represents a cell, and the *k* of an M-D cell pair is assigned to the daughter cell. **(e)** The logistic modelling for the hatching phenotype with the same settings in Fig. 4b. First, we only use cell-specific CCL features (similarly defined as cell-specific position features) as independent variables and conduct modelling in all embryos, obtaining the ROC curve and corresponding AUC (±standard deviation). To consider the potential basis from imbalanced hatched-lethal ratio, a shuffled model is also conducted by shuffling the hatching phenotypes. Then, we include the effect cells’ features identified based on position features in Fig. 4b but only make CCL features be subjected to feature selection in LASSO. This allows us to identify extra CCL features beyond the previously identified cell position features. **(f)** The distribution of effect cells in the cell lineage tree, with the newly identified cells by including CCL into the model. **(g)** Details of the high-dimensional CCL noise transmission model. Because the CCL information is incomplete for cells in the 10^th^ generation, a total of 345 M-D cell pairs from the 4^th^ to the 9^th^ generation are examined. **(h)** Summary of the coefficients in the *K* for CCL (*K*_CCL_), with standard boxplots shown. **(i)** The predicting performance of the full model for CCL and three nested models that consider only mother, only sister, or both mother and sister, respectively. **(j)** All eigenvalues of *K*_CCL_ have a negative real part. **(k)** The predicting performance of the full model for CCL and two nested models that block either mother or sister. **(l)** The eigenvalues of *K*_CCL_ in the nested models blocking mother. **(m)** The eigenvalues of *K*_CCL_ in the nested models blocking sister.

To assess the system stability against CCL noise, we followed Eq. (3) to build a high-dimensional model, shown in **Fig. 6g** where *L* represents the CCL noise of a cell in an embryo. Using the same method for analyzing Eq. (3) we obtained the matrix *K* and found it also sparse, with similar properties as the previous *K* matrices for position noise (**Fig. 6h-i and Fig. S23**). Importantly, the eigenvalues of the *K* also all have a negative real part (**Fig. 6j**), highlighting the stability of the nematode embryogenesis against CCL noise. The stability holds after blocking descendant cells (**Fig. S24**), or blocking sisters, but would vanish if mothers were blocked (**Fig. 6k-m**). All these results are parallel to the preceding findings in analyzing cell position noise.

## Discussion

Despite exceptions, previous studies on developmental canalization have focused on the regulations of molecular noises within a cell. In this study, we revealed in *C. elegans* ubiquitous mother-daughter negative-feedback regulations on the cell noises within an embryo. Because the factors modulating cell noises do not necessarily affect molecular noises, and the cell noises would still exist even if molecular noises are well controlled, our work suggests a new layer of noise regulations that are directly responsible for embryonic stability. Specifically, to achieve stable developmental end-products against perturbations, a multicellular organism seems to need two layers of regulations, one for suppressing molecular noises and the other for managing cell noises (**Fig. 7**). At the embryonic layer, the ubiquitous mother-daughter negative feedbacks revealed in this study can form continuous and comprehensive ‘canals’ for suppressing cell noise accumulation, wherein the estimated *k* and *d* quantitatively define the steepness and depth of the canals, respectively. We noted that, although the concept of canalization was introduced by Waddington 80 years ago, developmental canals have long remained to be a metaphor without concrete demonstration. Our work thus transforms, for the first time to the best of our knowledge, the metaphor into a mathematically formulated reality.

**Fig. 7.**
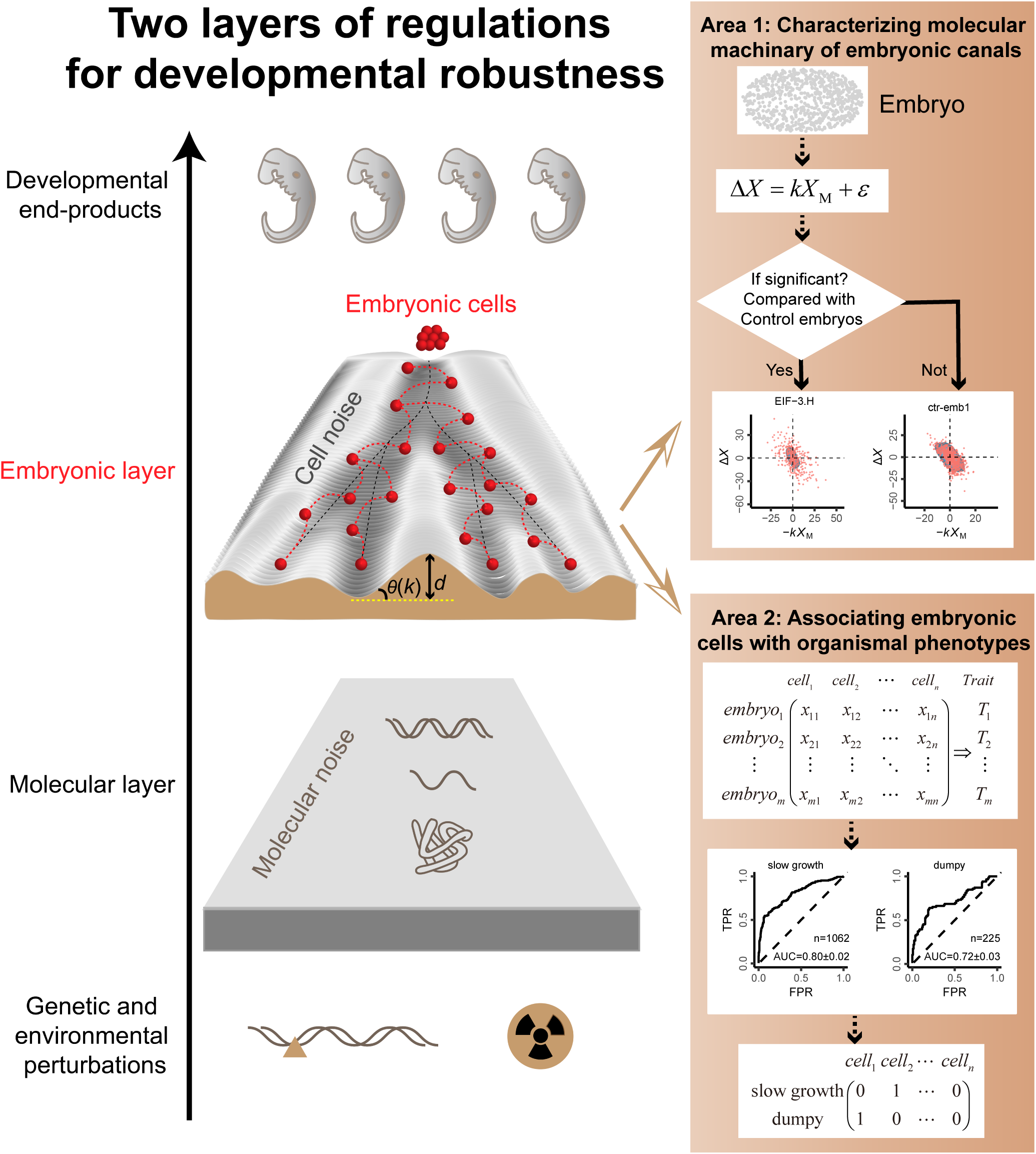
Two layers of regulations for developmental robustness. The embryonic layer of regulation, which is newly discovered and characterized in this study, directly contributes to developmental robustness. It employs continuous negative-feedback ‘canals’ to prevent cell noise accumulation, with two parameters, *k* and *d*, quantitatively defining the steepness and depth of the canals, respectively. The black dashed lines represent ideal cell trajectories along the canals, while the red dashed lines represent actual cell trajectories shaped by the canals. Notably, characterization of developmental canals at the embryonic layer suggests two future research areas: one for revealing the molecular machinery underlying the canals and the other for associating embryonic cells with organismal traits.

This study draws two immediate implications. First, there might be a novel molecular machinery underlying the developmental canals we revealed at the embryonic level (**Fig. 7**). Notably, characterization of the developmental canals provides a quantitative tool to identify the related genes. Specifically, for a given gene knockout/knockdown embryo we can test if the embryo fits the wild-type noise transmission models to assess the gene’s function. The RNAi knockdown embryos examined in this study are generally similar to the wild-type control embryos in terms of the noise suppression. However, some of the knockdowns may have significant whole-embryo defects relative to the wild-types, suggesting the perturbed genes are likely critical for maintaining the developmental canals. Among the RNAi knockdowns we identified 33 with statistically significant poorer performance than the wild types in suppressing the cell noises (**Table S8; Methods**). The perturbed genes, which are critical for maintaining the canals, are enriched in processes related to translation initiation and with phenotypes related to cell division timing (**Fig. 7 and Fig. S25**). Many of these genes have orthologs that are associated with developmental diseases in humans (**Table S8**); for example, the gene *LARS2* (a human ortholog to *lars-2* in *C. elegans*) is associated with premature ovarian insufficiency^46^. The identification of these genes marks a starting point for characterizing the molecular machinery responsible for the developmental canals in *C. elegans*. It is plausible that the to-be-characterized molecular machinery would be as vital as those for managing molecular noises, such as Hsp90, in shielding multicellular organisms from disturbances.

Second, this study suggests a strategy for associating embryonic cells with organismal phenotypes (**Fig.7**). We found the noise level of embryonic cells can accurately predict hatching lethality, thereby linking specific cells with an organismal trait. With additional phenotypic trait information available, we could conduct similar association studies to connect embryonic cells with organismal traits. This proposal is reminiscent of classical association studies in quantitative genetics that link genetic variants with traits. As such, we would understand better how embryogenesis per se shapes organismal traits. As a preliminary test, we assigned the phenotype of the RNAi embryos examined in this study by referring to the phenotypic traits reported in WormBase ParaSite (**Methods**). The required assumption is that the knockdown embryos of the same gene show the same organismal phenotype across the studies. We found the noise level of embryonic cells explains substantially the 20 assessed organismal traits such as “slow growth” and “dumpy” (**Fig. 7, Fig. S26 and Table S9**), highlighting the potentials of the proposed novel association study framework.

Nevertheless, there are serval caveats or limitations in this study. First, we considered cell position noises and cell cycle length noises separately in this study. Future studies could use a more sophisticated model that simultaneously includes spatial and temporal data to better characterize noise transmission. Also, considering certain deterministic non-linear model could be helpful^47^. Second, the between-cell regulations were characterized based on observational data. A perturbational method able to alter an individual cell’s position in an embryo would be desirable for further testing such regulations. However, currently there are no such methods available for *C. elegans* although optogenetic method can achieve targeted cell ablation^48^. Third, the nematode *C. elegans* is a simple multicellular organism with fixed developmental cell lineages. It remains unclear how the principles learned in this study would apply to more complex organisms. Finally, morphogen gradients^49^, cell-cell adhesion^50^ and other self-organization principles^51^ have also contributed to our understanding of embryogenesis. It’s intriguing how to integrate our discovery with these prior findings.

## Data and Code Availability

The raw data, processed data and codes for this study can be accessible via the following GitHub repository: https://github.com/Jianguo-Wang/canalizationENV. The spatial-temporal data of RNAi and control embryos can be obtained from our previous study^33^.

## Methods

### Single-cell-resolution Spatial-temporal Tracking of *C. elegans* Embryos

#### Environmentally-perturbed Embryos

A total of 120 embryos were generated through experiments involving 8 types of environments with 40 conditions, each condition consisting of 3 embryos. Under different temperature conditions (16℃, 18℃, 20℃, 22℃, 24℃) during the 2-4 cell stage, all embryos hatched normally. Similarly, under different heat shock time conditions (10 min, 15 min, 20 min, 25 min, 30 min) at 30℃ during the 2-4 cell stage, all embryos hatched normally. Culturing nematodes with different bacterial food types (HB101, OP50, BW25113, HT115, and DA1877) from the L1 larval stage for two continuous generations, resulted in all embryos hatching normally. Additionally, culturing nematodes with different stearic acid (SA) concentrations (50 μg/mL, 100 μg/mL, 200 μg/mL, 400 μg/mL, 600 μg/mL) and different glucose (GLU) concentrations (50 mM, 100 mM, 200 mM, 400 mM, 600 mM) from the L1 larval stage led to all embryos hatching normally, except for the 600 mM glucose concentration which resulted in lethality. Similarly, different sodium chloride (NaCl) concentrations (20 mM, 40 mM, 60 mM, 80 mM, 100 mM) during culturing produced normally hatched embryos. Treating nematodes with different iron chloride (Fe) concentrations (1 mM, 2 mM, 4 mM, 6 mM, 8 mM) at the L1 stage and under the 8 mM condition at the L4 stage resulted in all embryos hatching normally. However, treating nematodes with different paraquat (PQ) concentrations (1 mM, 2 mM, 4 mM, 6 mM, 8 mM) at the L4 stage caused lethality in all embryos. Out of the Env embryos, 11 (corresponding to 5 conditions) exhibited embryonic lethal phenotypes at the hatching stage, which were successfully traced until the 350-cell stage.

#### RNAi-perturbed and Control Embryos

In our previous study^33^, we carried out experiments on RNAi and control embryos in *C. elegans*, cultured under standard laboratory conditions at 21°C, unless specified otherwise. We focused on protein-coding genes with human orthologs, located on chromosome I, extracted from the Ensembl database. Expression data for these genes at the 4-cell stage, proliferating and gastrulating embryos, were procured from the EBI Expression Atlas, produced by the modENCODE project. Genes expressed at a transcript per million (TPM) > 5 at any stage were considered, which totaled 922 genes. We verified the accuracy of inserts for 752 expressed genes by sequencing available RNAi clones from the Ahringer RNAi library. RNAi treatments were administered using the feeding method, with slight modifications. Synchronized L1 larvae worms (20-25 worms per experiment) were moved to RNAi plates, containing bacteria that expressed double-stranded RNA specific to target genes. The worms were subjected to RNAi for 36-48 hours at 21°C, with experiments conducted using embryos from the treated worms. In instances where severe defects such as larval lethality or sterility made it impossible to collect embryos, a weak RNAi treatment was applied to L3-L4 stage larvae prior to the experiments. The 105 control embryos, which were fed with bacteria carrying the empty L4440 RNAi vector, all normally hatched. Out of the RNAi embryos, 98 (corresponding to 63 genes) exhibited embryonic lethal phenotypes at the hatching stage, which were successfully traced until the 350-cell stage.

#### 3D time-lapse Microscopic Imaging

As in our previous study^33^, gravid young adult worms (3-4) were dissected in egg buffer on a multi-test slide. Embryos before the 4-cell stage were selected and transferred onto a coverslip with polystyrene microspheres. The embryos were arranged and covered with a second coverslip before sealing the slide with Vaseline. For imaging, we employed a spinning disk confocal microscope (Revolution XD) with two channels (488 nm for GFP and 561 nm for mCherry). Imaging was performed using MetaMorph software under a 60x oil objective at 21°C. We acquired images for 4.75-6 hours with 75-second intervals, scanning 30 Z planes with 1-µm spacing for the mCherry and GFP channels. We optimized imaging parameters to minimize photobleaching and phototoxicity while maintaining a high signal-to-noise ratio. After imaging, we determined the hatching status of the embryos 24 hours later. Among the imaged wild-type embryos (304/308), 98.7% hatched normally without apparent morphological abnormalities. This hatching rate is comparable to that observed under standard NGM culture conditions, indicating minimal disruption of embryogenesis by the live imaging procedure used in this study.

#### Cell Lineage Reconstruction and Acquisition of Spatial-temporal Features

As in our previous study^33^, 3D time-lapse image series were processed with StarryNite software to identify and trace nuclei, enabling cell lineage reconstruction. Nuclei were automatically recognized from the histone::mCherry fluorescence in each time point’s 3D images, using hybrid blob-slice detection algorithms. The recognized nuclei were then traced over time to construct the cell lineage using semilocal neighborhood-based algorithms. Each traced cell was assigned a unique name based on Sulston’s naming rules. Cell lineages were traced until the 350-cell stage, unless severe developmental defects occurred due to gene knockdown. The digitized lineage data provided single-cell-resolution spatial-temporal features, including cell birth and division times and 3D cell positions at each time point. Three types of lineaging errors were manually detected and corrected: early termination, excessive length, and excrescent lineage branching. To ensure accuracy, the identification and tracing results underwent multiple rounds of manual curation using AceTree software. The accuracy of cell lineage results was assessed by examining 378 randomly selected traced terminal cells from 50 control embryos. Only one cell tracing error was identified, resulting in a cumulative accuracy of 99.9%. Residual errors, although present, they did not significantly impact the overall conclusions of this study.

### Alignment of Cell Cycle Lengths Among Embryos

The objective of this study is to quantitatively characterize developmental canals. This involves modeling relationships between mother and daughter cells, cell features in relation to the organismal phenotype, as well as high-dimensional interactions among cells. Consequently, proper alignment of cell features is crucial. We first examine the cell cycle length (CCL), which is defined as the time interval between two consecutive cell divisions.

#### Scaling

We notice that there are variations in the overall pace of development. Some embryos develop at a slower rate (with overall larger CCLs) or a faster rate (with overall smaller CCLs) than others. For each embryo, we quantify this variation by estimating a scaling factor (SF), which is determined as a relative scaling value to our reference embryo, the ctr-emb58. For instance, if an embryo’s cells have a two-fold CCL compared to the reference embryo, the scaling factor for the CCL of the focal embryo is 2. We derive the scaling factors for each coordinate of every embryo by performing a passing-origin orthogonal regression between the reference’s coordinate (independent variable) and the focal embryo’s coordinate (dependent variable). The slope represents the estimated scaling factor. By multiplying the raw CCLs by the reciprocal of the SF (1/SF), we obtain the scaled CCLs for every embryo.

#### Evaluation of Alignment Quality

To ensure the quality of alignment, we use the Identity Score (IS). The IS is defined as the variance proportion along the diagonal line when an embryo’s CCLs are plotted against those of the reference embryo. This score helps us understand how well the CCLs of the embryos align with the reference. Generally, a larger IS suggests a higher quality of alignment. However, a smaller IS may also result from an increased noise level in the considered embryo.

#### Impact of Reference Selection

We also evaluate the impact of the choice of reference on the SFs. This is done by comparing SFs based on ctr-emb58 to those based on the average CCL features among all control embryos. The good fitting between them by a line passing the origin indicates that the CCL features based on different references can be equivalently transformed by a fold change, suggesting a minimal effect from the reference selection.

### Spatial Coordinate Alignment via Translation-Rotation-Scaling Procedure

Time-lapse microscopy outputs three raw coordinates: *x*, *y*, and *z*. However, direct comparisons among embryos using these raw coordinates may not be unsuitable, as each embryo’s positioning within the culture and the scaling across all three axes can vary. To enable consistent comparisons across embryos, we applied a translation-rotation-scaling approach, as detailed below:

#### Translation

We first calculated each embryo’s center of gravity by averaging the three raw coordinates. We then derived the translated coordinates by subtracting the center of gravity from the corresponding raw coordinate.

#### Rotation

Assuming comparable body axes among embryos, each cell could be visualized as a point in a three-dimensional space, using the three body axes as Cartesian axes. In this framework, raw coordinates are perceived as a linear transformation of the coordinates that use the three body axes as Cartesian axes. To recover the three body axes, we use a Principal Component Analysis (PCA) method. In this method, the first, second, and third components correspond to the Anterior-Posterior (A-P) axis, Left-Right (L-R) axis, and Dorsal-Ventral (D-V) axis, respectively. For atypical embryos with unconventional body axes - for example, a shorter L-R axis than D-V axis - we designated a reference embryo (ctr-emb58) with standard body axes. The PCA-based linear transformation ensured the independence of the rotated coordinates.

#### Scaling

Following translation and rotation, variations in the overall size of the three axes among embryos were evident. We quantified these variations with Scaling Factors (SFs), defined and estimated in the same manner as the SF of CCL. The embryo ctr-emb58 served as the reference. We then derived the scaled coordinates from the rotated coordinates with these SFs.

#### Evaluation of Alignment Quality

We assessed the alignment quality from two perspectives. First, we computed the Identity Score (IS), defined and estimated similarly to the IS of CCL. Second, we evaluated the degree to which each coordinate (*x*, *y*, and *z*) of an embryo corresponded to the same coordinate of the reference embryo, while being mostly independent of the other two coordinates of the reference embryo. This secondary evaluation aimed to assess any misalignments or false designations.

#### Impact of Reference Selection

We evaluated the impact of reference selection on the alignment of spatial coordinates by recalculating the Scaling Factors (SFs) based on the average of all control embryos. We found that these align well with those obtained based on ctr-emb58.

#### Alignment of Coordinates at Cell Division

Our study tracked each embryo’s entire lifecycle from birth to division. Although our analysis primarily focused on position features at birth, considering that the negative-feedback effect could continuously operate from cell birth to division, aligning the position features at division was also useful. To maintain a consistent Cartesian coordinate system, we used the translation matrices, rotation matrices, and scaling factors obtained during the translation-rotation-scaling procedure applied to the position features at birth, to perform transformations on the position features at division.

### Mother-daughter Negative Feedback Model

#### Negative Feedback Model for Position Features

The negative feedback model for position features utilized in this study aligns with previous studies^39,40^, which has been described in detail in section “Modeling Cell Noise Transmission Along Developmental Lineages” of the main text. Here, we will only provide some necessary supplementary explanations. A coefficient *a* < 1 means if mother deviates from its expected position, the deviation will be suppressed in the daughter. Since *k* = *a* - 1, a *k* < 0 (equivalent to *a* < 1) serves as the criterion to assess mother-daughter negative feedback. The significance of *k* in Eq. (2) is determined by that of *a* in Eq. (1).

#### Negative Feedback Model for Cell Cycle Length (CCL)

We model the relationship between the CCLs of a daughter cell (*l*_D_) and those of its mother cell (*l*_M_) as follows:

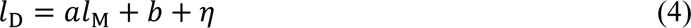

which is equivalent to:

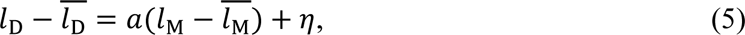

where *l* represents a cell’s CCL and 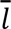 is the average CCL across embryos. Then, 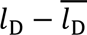 and 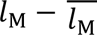 represent the noise of the daughter and that of the mother. Due to differing CCL expectations in embryonic cells, we further normalized the noise in Eq. (5) by the average of the daughter and that of the mother, respectively, as:

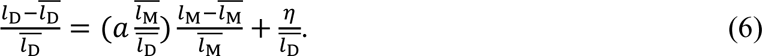

Then, 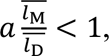 which is equivalent to 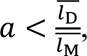 implies that when the mother deviates for a certain proportion, the daughter tends to inherit a smaller proportion of the deviation, aligning with a previous study^40^. By transforming Eq. (6) into a unified form for different mother-daughter pairs, we get:

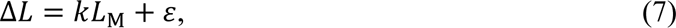

where 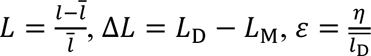 and 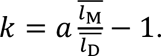 Therefore, *k* < 0, equivalent to 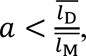 is used to judge mother-daughter negative feedback. The significance of *k* in Eq. (7) is evaluated in the context of the slope 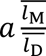 in Eq. (6) or the slope *a* in Eqs. (4, 5).

#### Continuous Negative Feedback Model Considering Position Features of Mother and Daughter at both Birth and Division

The model settings are largely the same, with one exception. We need to replace the *X*_M_ at birth with *X*_M_ at division when we consider the noise transmission from the division of mother to the birth of daughter (X_Mdivision_ − X_Dbirth_). Similarly, the other two situations ( X_Mdivision_ − X_Ddivision_ and X_Mbirth_ − X_Ddivision_) can be derived. Our comparison among the four combinations revealed that a longer interval between a mother’s stage and a daughter’s stage corresponds to stronger negative feedback. Despite potential slight overestimation of the *k* values for X_Mdivision_ − X_Ddivision_ and X_Mbirth_ − X_Ddivision_ due to incomplete records of cell division information for cells in the final generation, the relative interval length for the four combinations is maintained.

### Analysis of Potential Confounding Factors

#### Random Mother-Daughter Cell Pair

To rule out potential data structure issues that could cause consistently negative *k* values, we created pseudo mother-daughter (M-D) pairs. This was accomplished by shuffling the corresponding daughter cells in relation to the mother cells.

#### Scaling

The scaling process could either stretch or compress raw coordinates. We examined whether negative *k* values were an artifact of this process. We grouped the embryos into a series of ranges based on their scaling factors (SFs), using the median SF as the midpoint (ranging from 10% to 100%). A narrow range around the median SF represents embryos with similar scales and hence, less impacted by the scaling process. Within these ranges, we observed that the *k* values, computed based on each range of embryos, predominantly remained negative. This suggests that the negative *k* values are not merely a byproduct of the scaling process.

#### Outlier Effect (Low Alignment Quality)

Extreme outliers could skew the estimation of the slope in ordinary linear regression. In this study, this issue could primarily arise from embryos with poor alignment quality. We filtered out embryos with relatively lower alignment quality, specifically those with an IS ≤ 0.9, leading to the exclusion of only a few dozen embryos. We then re-estimated the *k* values. The results indicated that while the exclusion of outlier embryos could marginally reduce the *k* values, the re-estimated *k* values maintained a strong correlation with the *k* values derived from all embryos.

#### Dilution Effect

When an independent variable contains measurement errors, the estimated slope between the independent variable and dependent variables becomes attenuated towards the independent variable. This phenomenon is known as the dilution effect^52^. In this context, the measurement error-adjusted slope (*a*^adjusted^) and the observed slope (*a*^observed^) between the mother cell and daughter cell are represented by 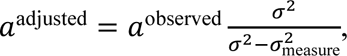 where *σ*^2^ is the mother cell’s variance and 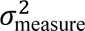 is the mother cell’s measurement error variance (MEV). In the study, MEV is determined by the smallest unit of measurement (0.25 min for CCL, 1 pixel for raw *x* and *y* coordinates and 4.5 pixels for raw *z* coordinates). Thus, each cell feature follows a uniform distribution U (O - *u*, O + *u*), where U stands for uniform distribution, O represents the observed value, and *u* equals 0.25 min for CCL, half of 1 pixel for raw *x* and *y* coordinates, and half of 4.5 pixels for raw *z* coordinates. Consequently, each cell feature’s MEV in a given embryo is calculated as *W* = 4*u*^2^/12, meaning each cell in the same embryo has identical MEV for a specific feature. As rotated coordinates are a linear combination of raw coordinates, a cell’s MEV for rotated *x*, *y* and *z* coordinates within a specific embryo is computed as 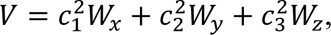 where *c*_1_, *c*_2_ and *c*_3_ represent the loadings of a rotated coordinate on the raw *x*, *y* and *z*. After scaling, a cell’s MEV for scaled *x*, *y* and *z* coordinates in a specific embryo is estimated as, *V* / SF^2^, where SF is the scaling factor of a rotated coordinate in *i*^th^ embryo. The MEV of a specific feature across all embryos is then calculated as the average of the MEVs among embryos, expressed as, 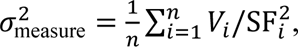 where *n* equals the number of embryos, *i* signifies the *i*^th^ embryo, *V_i_* represents the *i*^th^ embryo’s MEV, and SF*_i_* denotes the scaling factor of the feature in the *i*^th^ embryo. Finally, for the dilution effect-adjusted negative feedback for a mother-daughter cell pair is calculated as, 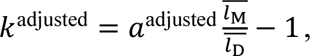 referring to the relationship between *k* and *a* in the previously described mother-daughter negative-feedback model. Our results clearly indicate that the dilution effect has a negligible contribution to the *k* values.

#### Tracking Error

When a tracking error is introduced into a specific generation, the error rate is computed as the proportion of misidentified cells to the total cell count. For example, if we introduce a 5% error rate in the 4^th^ generation for the *x* coordinate, given that the 4^th^ generation is our starting point for measurements, a single error in this generation can propagate errors into subsequent generations. In this scenario, we randomly select 5% of the mother cells in the 4^th^ generation across all embryos and interchange the two daughter cells for each of these selected mother cells, as well as their subsequent generations. This manipulation corresponds to an error rate of 5%. Subsequently, we re-estimate *k*_x_ based on the data with the introduced error. The resulting accuracy is calculated as the square of Pearson’s *R* between the original *k*_x_ (calculated based on the assumed error-free measured data) and the re-estimated *k*_x_, which now includes a 5% error rate. Introducing an error rate of up to 10% into each generation (less than 1% for the RNAi data, based on the manual examination in the previous study^33^), we observed consistently high accuracy. This suggests that the *k* values display significant robustness against tracking errors.

#### Alignment Procedure

To recover the true three body axes, we implemented a translation-rotation-scaling alignment procedure. This procedure is inevitably affected by cell noise, particularly because the PCA-based rotation process itself is a variance-based method. To assess the impact of this alignment procedure on the estimation of *k* values, we conducted a simulation. In this simulation, we set theoretical *k* values and subjected the embryos to a reverse process, namely, scaling-rotation-translation. We then calculated the observed *k* values following the same alignment procedure used in our study. We used the square of Pearson’s *R* to evaluate the robustness of observed *k* values compared to the theoretical *k* values. Specifically, we retained the cell noises in the 4^th^ generation and the *k* values for the 4^th^ to 5^th^ cell pairs unchanged. We then shuffled the residuals in Eqs. (1, 2) to generate the simulated cell noises in the 5^th^ generation. Subsequently, we generated the simulated cell noises in the 6^th^ generation based on the simulated cell noises in the 5^th^ generation and the *k* values for the 5^th^ to 6^th^ cell pairs. The cell noises in the 7^th^ to 10^th^ generations were generated in a similar manner. In accordance, we generated the same number of simulated embryos in which the theoretical negative feedback or *k* for each cell pair was set as the obtained *k* values in our study. We applied the reverse procedure (scaling-rotation-translation) to these simulated embryos and obtained simulated observed embryos. We then applied the same procedure (translation-rotation-scaling) used in the real embryos to estimate *k*_x_, *k*_y_, and *k*_z_ in the simulated observed embryos. Finally, we compared the theoretical *k* and observed *k* in simulated embryos and found them to be highly correlated. Since the same translation-rotation-scaling procedure is applied to all cells in the same embryo, both mother and daughter cells, we do not anticipate that the *k* adjusted by the procedure would exhibit the same directional deviation as that adjusted by the dilution effect.

#### The σ_D_/σ_M_<1 Issue

In this study, we estimate the negative feedback *k* values through ordinary linear regression, where the slope (*a*) and Pearson’s *R* between dependent and independent variables satisfy *a* = (σ_D_/σ_M_)×*R*. Here, σ_D_ and σ_M_ denote the standard deviation of daughter and mother cells, respectively. Given that *R* ranges from -1 to 1, it follows that *a* < σ_D_/σ_M_. When σ_D_/σ_M_ < 1 is a common property of mother-daughter cell pairs, it would always be true that *a* < 1, and consequently, *k* = *a* – 1 < 0. In this scenario, validating negative feedback using negative *k* values becomes inappropriate. Furthermore, if σ_D_/σ_M_ could explain the main variance of *k* values, this would suggest that the derived *k* values primarily originate from the upper limit of σ_D_/σ_M_, and thus may lack actual significance. For cell cycle length (CCL), we normalize raw CCL noise relative to the average CCL of each cell, which could inadvertently introduce a σ_D_/σ_M_ < 1 issue due to the consistently longer CCL of daughter cells relative to mother cells. The σ_D_/σ_M_ < 1 problem for CCL can also be demonstrated as a smaller coefficient of variation (CV) for the cell noise of daughter cells compared to that of corresponding mother cells. However, our results demonstrate that σ_D_ and σ_M_ are comparable, and σ_D_/σ_M_ only accounts for a portion of the variance in *k* values, indicating that our findings are not artifacts of this potential issue.

### Null Model of Noise Transmission

#### Calculation of Expected Cell Noise Variance

For a lineage comprised of seven cells, such as ABal-ABala-ABalaa-ABalaaa-ABalaaaa-ABalaaaal-ABalaaaala, we calculate both the observed and expected noise variance for each cell among embryos. For the observed noise variance, we calculate the variance of each cell’s noise across embryos. For the expected noise variance, we first acquire the cell-specific noise for the cells in the 5^th^ to 10^th^ generations within the lineage using Eqs. (1, 2), denoted by *ε*. Subsequently, we calculate the variance of this cell-specific noise among embryos for each cell. Following this, for a cell in the 5^th^ to 10^th^ generations, the expected noise variance is calculated as the sum of the cell-specific noise variance and the expected noise variance of its mother cell. For a cell in the 4^th^ generation, the expected variance is equivalent to its observed noise variance, as the 4^th^ generation is our initial recorded generation.

#### Generation of Expected Cell Positions within an Embryo

To generate expected cell positions, we first obtained the daughter-specific noise, *ε*, based on Eqs. (1, 2). Then, for a given embryo, the expected cell noises of daughter cells were generated by adding the expected cell noises of mother cells and the daughter-specific noises. The only exception is for the cells in the 4^th^ generation, whose expected noises are set to be the same as the observed noises. This suggests that the noise originating from mother cells is fully inherited by daughter cells. Finally, the expected cell positions were generated by adding the expected cell noises to the averages of cell positions observed in this study.

### Identification of Perturbed Cells

We first extract the *k* values and their corresponding standard errors using ordinary linear regression. Focusing on a subset of embryos - including control embryos and those exposed to specific perturbations - we re-estimate the *k* values and their standard errors. We then perform a pairwise two-tailed t-test to assess statistical significance. This process allows us to identify which cells, under specific perturbations, show significantly different *k* values compared to those derived from control embryos. The Benjamini-Hochberg method is used to generate the adjusted p-values.

### Tissue Enrichment Analysis of Cells

In a previous study^36^, differentiated tissue types or instances of apoptosis were identified for the cells in the final generations. We use this information to determine the tissue destinies for each cell examined in our study. If all descendants of a focal cell in the final generations belong to the same tissue type, we assign that tissue type to the focal cell, indicating that its fate aligns with that tissue type. If some descendants of a cell are destined for apoptosis in the final generations, but all remaining descendants belong to the same tissue type, we also assign that tissue type to the focal cell as its tissue fate. However, if some descendants in the final generations are unidentified in terms of their differentiated tissue type or apoptosis, or if these descendants belong to more than one tissue type, we classify the focal cell as having an undefined tissue fate. To evaluate significant tissue type enrichment within a set of cells - for instance, the perturbed cells identified based on *k* value comparisons - we perform a one-tailed binomial test, generating a p-value for each tissue type. Cells without an assigned tissue fate are not included in this analysis.

### Prediction of Embryonic Lethality

#### Model Setting

Successful embryogenesis in *C. elegans* hinges on the exact regulation of Cell Cycle Length (CCL) and accurate cell positioning during development. This insight presents the possibility of cell noise as a predictor of organismal phenotype. In this study, we recorded the hatching phenotype, namely, hatched or lethal, for each embryo at the hatching stage. We used a machine learning framework to develop a logistic regression model. This model utilizes cell-specific noise, which is independent between mother and daughter cells, thus allowing for the identification of potential effects from mother cells beyond those from daughter cells. Concurrently, the application of the LASSO method allows us to pinpoint a core set of cells that primarily contribute to embryonic lethality, which implies that the noise within these cells is instrumental in monitoring the normal hatching of an embryo, leaving minimal variance for the remaining cells which are then assigned a zero coefficient. Such cells, bearing non-zero coefficients in the learned logistic regression model, are designated as "effect cells."

#### Modelling and Evaluation

The predictability of cell-specific noise towards embryonic lethality is initially evaluated using a 2:1 training-testing split amongst all embryos, along with a ten-fold cross-validation within the training set. To ensure the independence, embryos of the same RNAi are mutually exclusively divided into training or testing sets. Subsequently, a ten-fold cross-validation utilizing all embryos is employed to construct a final model that identifies the effect cells. To assess the potential impact of an imbalanced hatched-lethal ratio, we shuffle the hatching state of embryos, building a corresponding shuffled model within the same learning framework. Model performance is assessed by generating the Area Under the Curve (AUC) and its corresponding standard error. Moreover, to test the prediction performance of early cells, we restrict the predictors by excluding cells after a given generation. It’s noteworthy that the "effect cells" represent the distribution of the predictive effects of noise within the lineage tree. However, their identification isn’t exclusive due to the correlated noise among sister cells. The R package "glmnet" facilitates the modeling process.

#### Combining Position and CCL Features

Given that we have identified effect cells’ features in the modelling of hatching phenotype based on position features, we combined the cell-specific CCL features with the identified position features to evaluate the extra contribution of CCLs. To avoid the identified position features from being subjected to feature selection in LASSO, we only allow the CCL features to be subjected to LASSO’s feature selection.

### Definition and Estimation of Noise Tolerance Level

Given the significant influence of cell noise on embryonic lethality, we posit that each cell has a specific noise tolerance threshold. If a cell’s noise exceeds this threshold, the probability of embryonic death dramatically increases. In this context, we define a cell’s noise tolerance level (*d*) as the level of cell noise corresponding to a 50% probability of embryonic lethality, as determined by logistic regression. The model between a cell feature’s absolute noise (|*X|*) and hatching phenotype is modeled by logistic regression logit(P) = E(*β*_0_+*β*_1_|*X|*), where P represents the probability of a lethal hatching phenotype, and E represents expectation. As such, *d* = -*β*_0_/*β*_1_, corresponding to 50% probability of hatching lethality. In cases where an extremely high noise tolerance level is obtained, or when the probability of embryonic lethality marginally decreases with increasing noise—indicating that the cell noise observed across embryos for a given cell does not predict embryonic lethality—we resort to using the cell’s maximum observed noise level among all embryos as a proxy for its noise tolerance level. For position features, we use the absolute form of calculated noise thresholds. However, for CCL, we use the relative percentage, namely, the absolute noise thresholds divided by the average CCL of each cell among scaled embryos. This is because the CCL noise we used to estimate *k*_CCL_ and to model hatching phenotype, is normalized to the average CCL of each cell. This adjustment makes the CCL noise tolerance level of cells across different generations comparable.

### High-Dimensional Stability Analysis of Embryonic Cell System

We examine the stability of the entire embryonic cell system in a high-dimensional space from two perspectives: mother-daughter stability across two cell cycles and birth-division stability within a single cell cycle. This analysis is performed for the *x*, *y*, and *z* coordinates respectively.

#### Mother-Daughter Stability across Two Cell Cycles

We illustrate the concept using the x-coordinate, but it applies to the y and z coordinates as well. We expand the mother-daughter (M-D) negative feedback model presented in Eq. (2) into a high-dimensional model expressed as Eq. (3), which encompasses all M-D pairs simultaneously. In this high-dimensional model, we investigate the stability of daughter cell noises 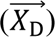 in relation to mother cell noises 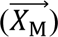 by examining the sign of the real parts of eigenvalues of matrix *K*. The element *K* (*i*, *j*) characterizes the effect of the *j*^th^ mother’s noise on the noise difference of *i*^th^ daughter cell from its mother’s noise.

#### Birth-Division Stability within a Single Cell Cycle

Again, we use *x* coordinate as an example, and the process can be similarly applied to *y* and *z* coordinates. We explore the stability of all cell noises at cell division 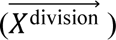 against those at cell birth 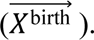 The resulting model is:

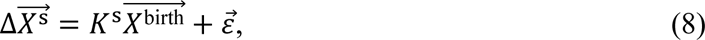

where 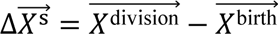 is a vector storing the noise difference of all cells between birth and division, and *K*^s^ is a coefficient matrix, with the item *K^s^*(*i*, *j*) representing the effect of *j*^th^ cell’s noise at birth on the noise difference of the *i*^th^ cell at division from its noise at birth.

#### Modelling

We estimate the coefficient matrices *K* in Eq. (3) and *K*^s^ in Eq. (8) using linear regression within a machine learning framework. We use the LASSO method in conjunction with ten-fold cross-validation to model each variable in 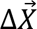 and 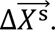 In estimating *K* in Eq. (3), we make two additional considerations. First, for mothers in Eq. (3) that have two daughters, we block one of the duplicated mothers in 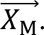 Second, for mother cells in the 5^th^ to 9^th^ generations that are also daughters of their own mothers, we block a mother cell itself in 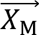 when modeling that mother. We refer to the full M-D high-dimensional model as the full model. In addition to this, we also consider several nested models. Specifically, for a focal daughter cell, we consider only mother variables, only sister variables, and both mother and sister variables in 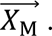 We also block only mother variables, and only sister variables in two other nested models. In modeling the birth-division stability within a single cell cycle, we take into account cells from the 4^th^ to 9^th^ generations. This is because the final 10^th^ generation contains incomplete data regarding cell positioning at division. For the modeling of mother-daughter stability across two cell cycles, we include cells from the 4^th^ to 10^th^ generations. The rationale behind this is that the cell position at birth is comprehensively recorded up until the 10^th^ generation. Moreover, the mother-daughter stability across two cell cycles for the Cell Cycle Length (CCL) is modeled in a manner akin to Eq. (3), considering cells from the 4^th^ to 9^th^ generations. In addition, we block the descendant cells for each daughter cell in 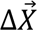 if they exist in the 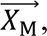 to exclude the potential impact of the descendant cells on the modelling.

#### Model Performance and Stability Analysis

Model performance is evaluated by the square of Pearson’s *R* between observed and predicted values for each item in 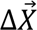 and 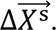 The stability of Eqs. (3, 8) is assessed by the eigenvalues of *K* and *K*^s^, respectively. If all eigenvalues of *K* and *K*^s^ have a negative real part, Eqs. (3, 8) are considered mathematically locally stable. For this analysis, Eqs. (3, 8) serve as approximations of 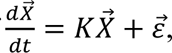 with dt approximated by a cell cycle.

### Effect of Cell Contact on High-Dimensional Stability

In a previous study^42^, the authors identified cell-cell contact information. For each daughter cell in 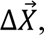 we gathered the mother cells in 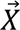 that are in contact with the focal daughter cell, as well as those that are not. We then extracted the coefficients in *K* for the two types of mother cells – those in contact and those not in contact with the focal daughter cell. The coefficients in *K* for each daughter cell in 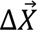 are respectively combined into two groups, “Contact” group and “No contact” group. We then computed the proportion of non-zero coefficients within each group. Furthermore, we compared the distribution of the absolute values of non-zero coefficients between the two groups. In this analysis, we focused on the coefficients excluding those corresponding to mother and sister effects due to their consistent dominating roles.

### Identification of Canal-Maintaining Genes

We first identify RNAi embryos whose CCL vectors significantly deviate from the mean CCL vector of 105 control embryos. Specifically, we calculate the Euclidean distance (D) between an embryo’s CCL vector and the mean CCL vector of the control embryos. By comparing the D values of the RNAi embryos with those of control embryos, we identify RNAi embryos with significantly larger D values at a significance level of 0.05 (Z-test, Bonferroni correction). From the embryos identified in the first step, we pinpoint RNAi embryos displaying substantial defects in mother-daughter (M-D) negative feedback. To achieve this, we calculate the residual vector in the M-D negative feedback model (Eqs. (1, 2)) for each M-D pair. We then compute the sum of squared residuals (SSE, Sum of Squares Error) for all cells in an embryo and normalize it by the sum of squared noises (SST, Sum of Squares Total) of these cells. We then identify embryos with significantly larger SSE/SST ratios than control embryos at a significance level of 0.05 (Z-test, Bonferroni correction). We similarly consider the *x*, *y*, and *z* coordinates. Finally, we identify candidate genes that maintain canalization, which correspond to the RNAi embryos identified through this two-step statistical testing process. The enrichment analysis for GO terms and phenotypes are conducted using the online tools of WormBase database. Human ortholog genes to these canalization-maintaining genes are obtained from DIOPT Ortholog Finder and gene-disease associations are abtained from DisGeNET accompanied by manual literature searching in PubMed.

### Prediction of Phenotypes from WormBase ParaSite

We collected the gene-phenotype annotation data from WormBase ParaSite 18. Given a phenotype, we assign the embryos of a gene’s RNAi as 0 if the gene’s ‘Qualifier’ is equal to ‘NOT’, and we assign the embryos of a gene’s RNAi as 1 if the gene’s ‘Qualifier’ is not equal to ‘NOT’. When a gene is not annotated in WormBase ParaSite 18, we assign a missing value for the embryos of the gene’s RNAi. After this, we obtained the 20 phenotypes each with at least 80 embryos assigned as 1. Finally, we apply the same logistic modelling for the 20 phenotypes as that used in modelling the hatching phenotype we recorded. Both the cell-specific position noise and CCL noise are used as predictors.

---

**Fig. S1.**
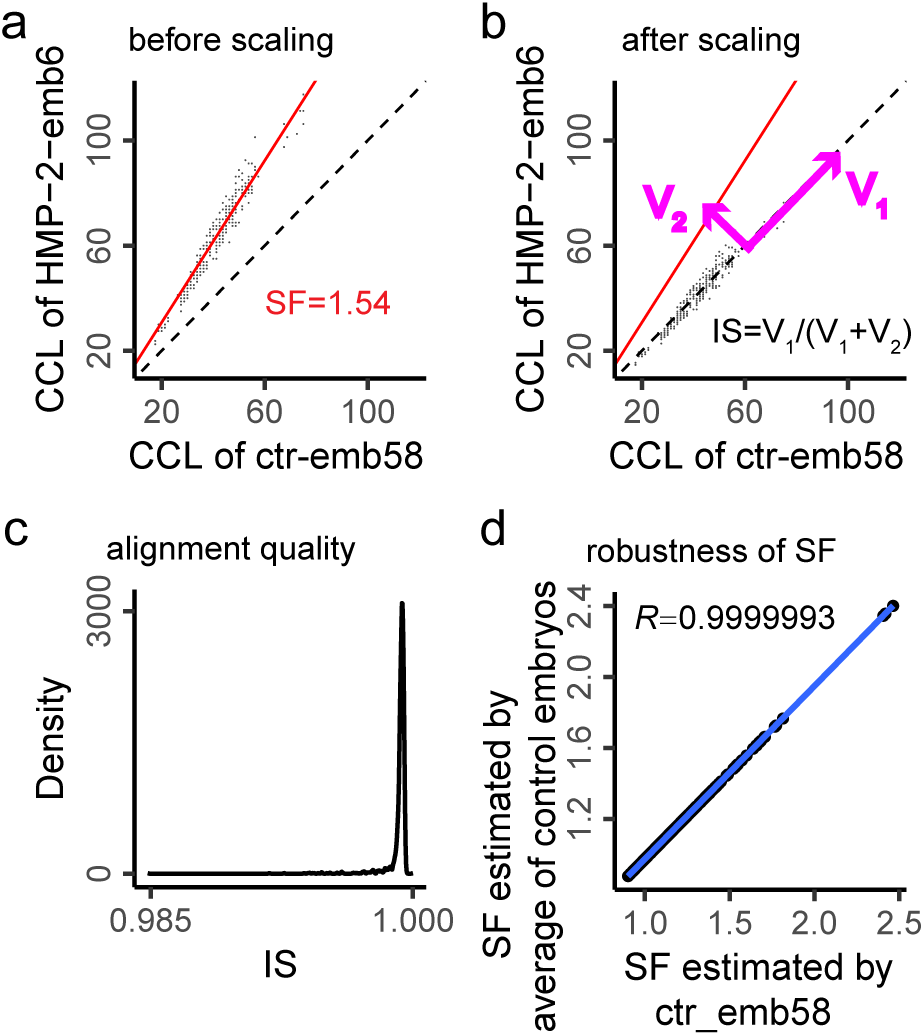
Alignment of Cell Cycle Length. **(a)** An example of an HMP-2 RNAi embryo (HMP-2-emb6). The Cell Cycle Lengths (CCLs) of HMP-2-emb6 are approximately 1.54 times those of the reference embryo (ctr-emb58), termed the Scaling Factor (SF). The SF is estimated by the slope of an orthogonal regression line passing through the origin, between the CCLs of HMP-2-emb6 and those of ctr-emb58. The dashed line represents the diagonal line. **(b)** Following SF estimation, the scaled CCLs are derived by dividing the raw CCLs by the SF. The scaled CCLs of HMP-2-emb6 are expected to align with the CCLs of ctr-emb58, meaning that data points should distribute along the diagonal line. The variance of these points can be decomposed into two components: one along and the other orthogonal to the diagonal line (V_1_ and V_2_). The proportion of variance along the diagonal line is defined as the Identity Score (IS), which is used to evaluate alignment quality. **(c)** The distribution of Identity Scores across all 2039 embryos. **(d)** Using the average CCLs of 105 control embryos as a reference, we re-estimated the SF for each embryo. The SFs obtained based on the two references can be well-fitted by a straight line passing through the origin (blue color), indicating that scaled CCLs based on two references can be approximately transformed by a fold change. This suggests that the choice of reference does not impact the analyses in this study. Ultimately, we chose a random embryo, ctr-emb58, as the reference instead of the average of control embryos. This is because the 105 embryos still have SFs (albeit with smaller variations than the perturbed embryos), which could introduce bias given that data with different scales do not follow the same distribution and are thus unsuitable for average calculation.

**Fig. S2.**
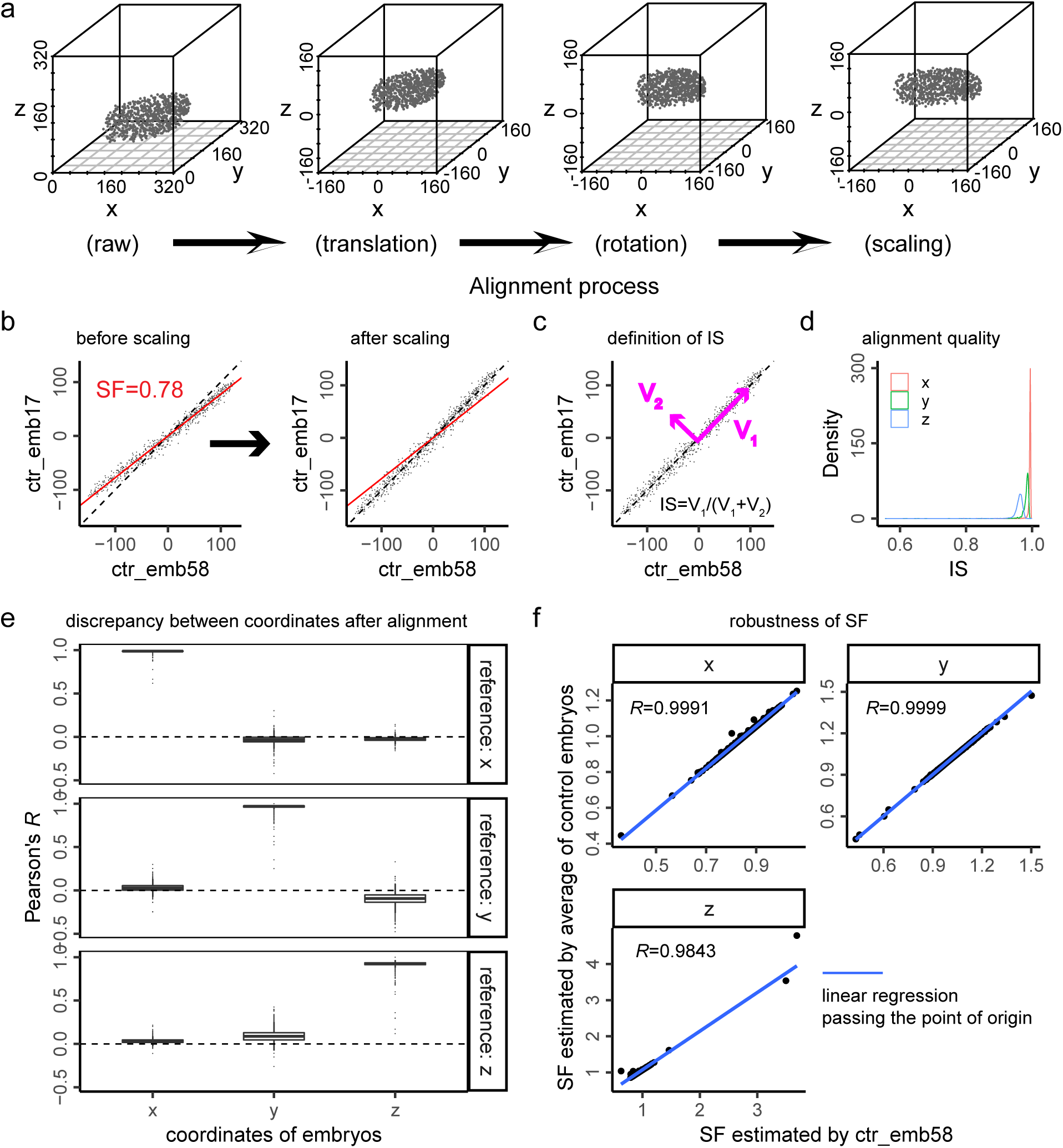
Alignment of Position Features. **(a)** The raw *x*, *y*, and *z* coordinates for each embryo undergo a sequential process of translation, rotation, and scaling to generate aligned coordinates across embryos. Each point in the three-dimensional plot represents a cell in an embryo. **(b)** The same scaling process employed for CCLs is used to scale the three position features. **(c)** Definition of the Scaling Factor (SF) for an embryo. **(d)** The distribution of SFs for embryos for *x*, *y*, and *z* coordinates, respectively. **(e)** Evaluation of coordinate misassignment in alignment. The horizontal axis represents *x*, *y*, and *z* coordinates of a focal embryo while the vertical axis represents the coordinates of the reference embryo. Pearson’s *R* is used to evaluate the similarity between any coordinate of a focal embryo and that of the reference embryo. **(f)** SFs remain robust when the average coordinates of control embryos are used as a reference.

**Fig. S3.**
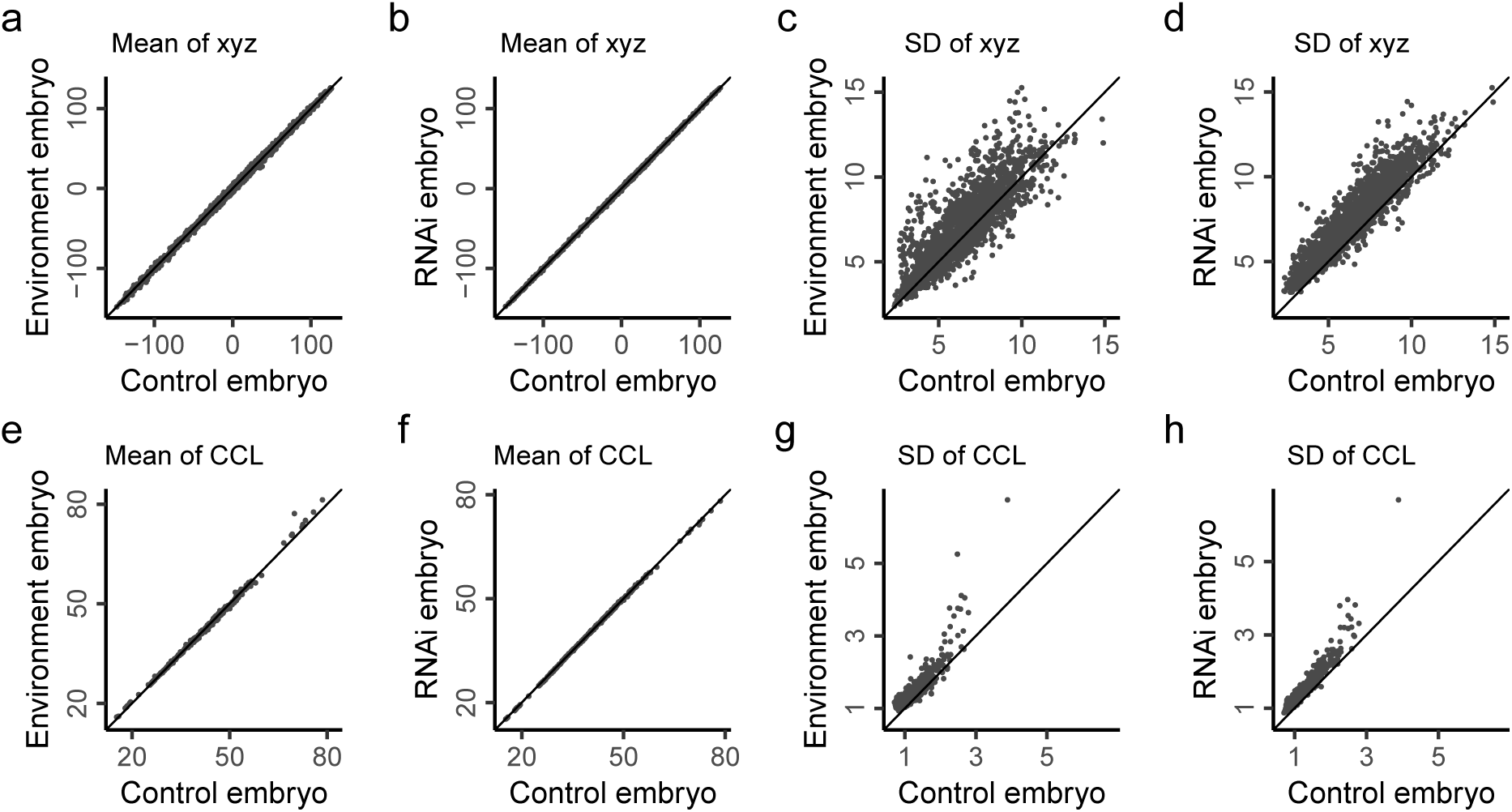
Comparison of Mean and Standard Deviation of Spatio-Temporal Features. In each panel, each point represents the mean or standard deviation (SD) of a particular spatio-temporal feature (after alignment), estimated based on two types of embryos. **(a-d)** Comparison for spatial features. **(e-h)** Comparison for CCLs. **(a-b and e-f)** Comparison for the mean of features. **(c-d and g-h)** Comparison for the standard deviation (SD) of features. **(a, c, e, and g)** Comparison between control and environment-perturbed embryos. **(b, d, f, and h)** Comparison between control and RNAi-perturbed embryos.

**Fig. S4.**
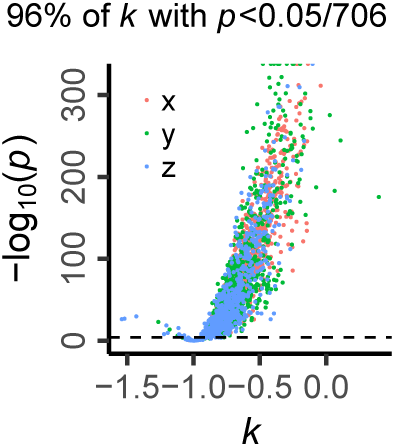
The Distribution of P-values for *k* Values of Spatial Coordinates. Note that the significance of the *k* value between a mother cell and the corresponding mother-daughter difference is determined by the significance of *a* in Eq. (1), since *k*=*a*−1. Therefore, we test the significance of *k* using the p-value (t-test) for the Pearson’s correlation between mother cell noise and daughter cell noise. The relationship between the *k* values and the p-values are shown for all mother-daughter cell pairs, for three coordinates respectively. The dashed line represents the threshold of *p*=0.05.

**Fig. S5.**
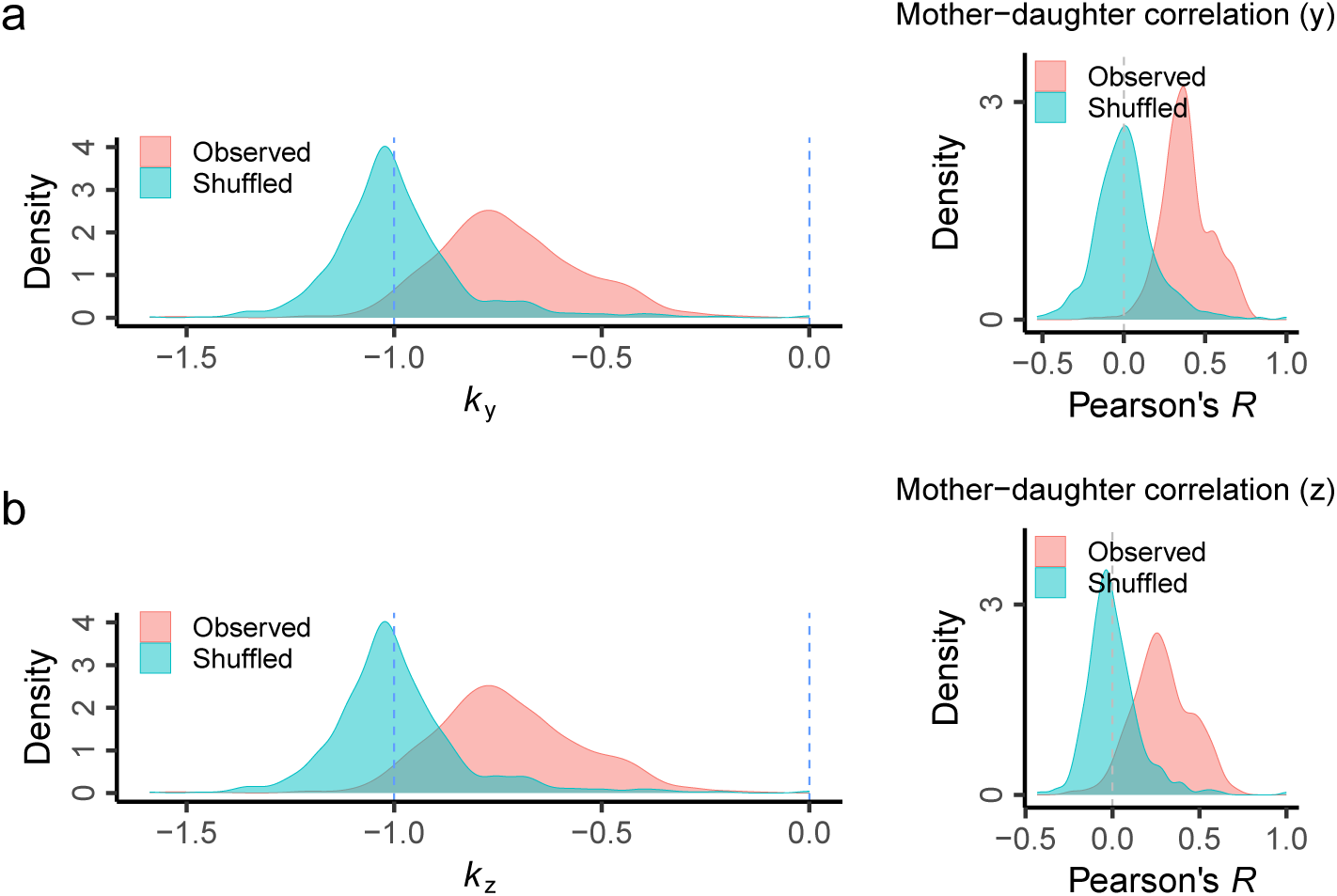
Comparison between Mother-Daughter and Random Cell Pairs. **(a-b)** Results for the *y* and *z* coordinates, respectively, corresponding to Fig. 2d. When a daughter cell fully inherits the noise of its mother cell, the expected *k* value is 0. When a daughter cell is fully independent of its mother cell, the expected *k* value is -1. When *k* lies between -1 and 0, a negative feedback regulation exists between mother and daughter cells. When *k* < -1, there are two possibilities. In the first scenario, the daughter cell still inherits the mother cell’s noise, but an over-negative feedback exists. In the second scenario, a negative correlation exists between the noises of mother and daughter cells. As expected in the situation of full independence, the shuffled *k* is distributed around -1. Most observed *k* values are distributed between -1 and 0, suggesting global negative feedbacks for mother-daughter noise inheritance. As expected in the case of full independence, the *R* value based on shuffled cell pairs is distributed around 0.

**Fig. S6.**
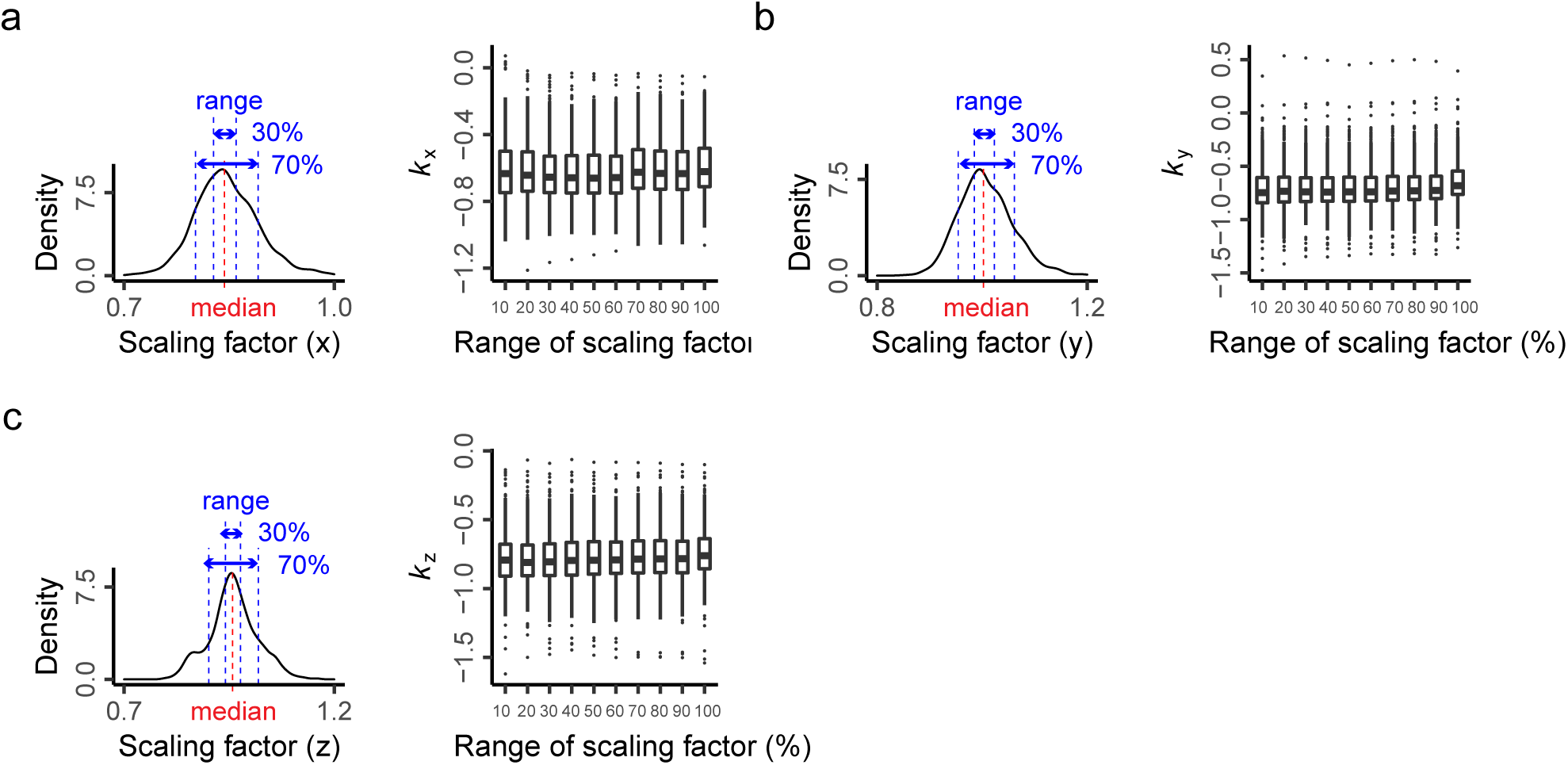
Robustness of *k* Against Scaling Factor. The left side of each panel shows the definition of the scaling range, which describes the percentage around the median of Scaling Factors (SFs). The distribution of SFs for the *x*, *y*, and *z* coordinates are shown, respectively. Outliers are not shown for clarity. The right side of each panel shows the estimation of *k*_x_, *k*_y_, and *k*_z_, respectively, given a range of scaling factors. A series of scaling factor ranges (from 10% to 100%) are considered.

**Fig. S7.**
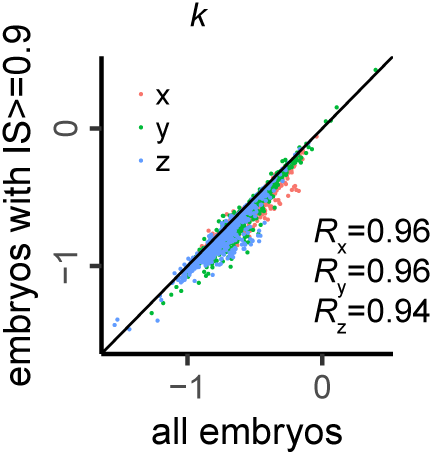
Estimation of *k* Values Using Embryos with High Alignment Quality. We define high alignment quality for an embryo if the Identity Score (IS) for all three coordinates of the embryo is greater than or equal to 0.9. This cutoff filters out 23 embryos. The re-estimated *k* values are plotted against those derived from all the embryos. Three coordinates are labeled in different colors. The diagonal line is shown for reference.

**Fig. S8.**
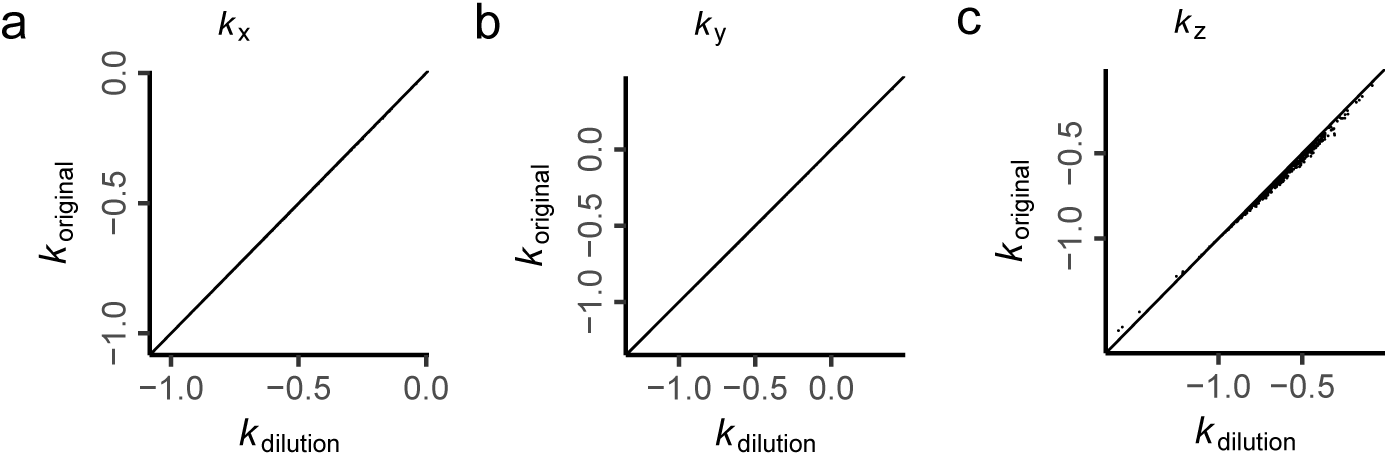
Dilution Effect of Measurement Precision on the Estimation of *k*. Each panel in this figure compares the *k* values estimated by considering the dilution effect from measurement precision with those estimated without considering this effect. The *k* values for the three coordinates are displayed separately, and a diagonal line is shown for reference in each panel.

**Fig. S9.**
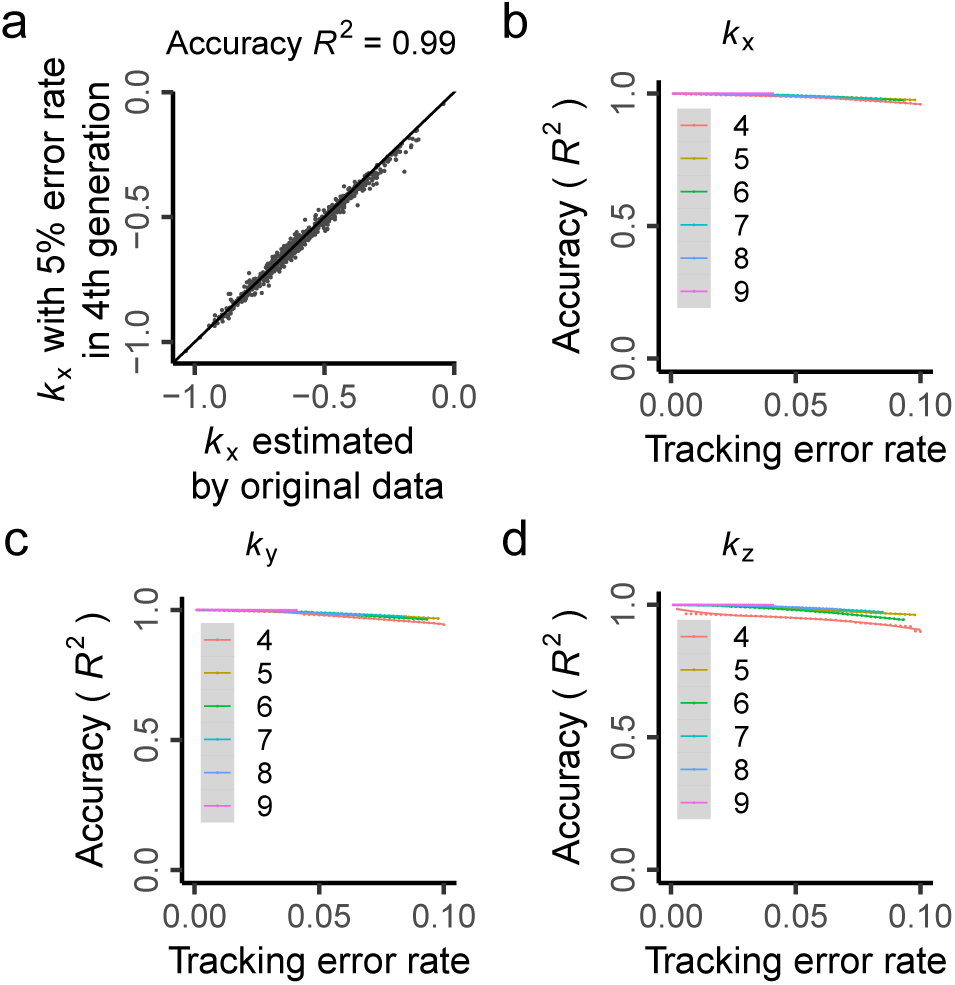
Impact of Tracking Error on the Estimation of *k*. Accuracy is defined as the square of Pearson’s *R* between the original *k* values and the *k* values re-estimated after introducing tracking error. When tracking error is introduced into a specific generation, the error rate is calculated as the ratio of the number of misidentified cells to the total cell count. **(a)** For example, we introduced a 5% error in the 4^th^ generation for the x coordinate. Specifically, we randomly selected 5% of mother cells in the 4^th^ generation across all embryos and interchanged the two daughter cells for each of these selected mother cells, as well as their descendants in subsequent generations. This corresponds to a 5% error rate. We then re-estimated *k*_x_ based on the data with this error. The *k*_x_ based on original data and the *k*_x_ estimated with a 5% error rate are highly correlated, resulting in an accuracy of *R*^2^=0.995. **(b-d)** Display the relationship between accuracy and error rate when tracking errors are introduced into different generations (mother generation: 4^th^-9^th^) for *x*, *y*, and *z* coordinates, respectively. The fitted lines are obtained by loess regression, and the grey bands represent the 95% confidence interval. As the previous study demonstrated, the tracking error rate in RNAi embryos is less than 1%.

**Fig. S10.**
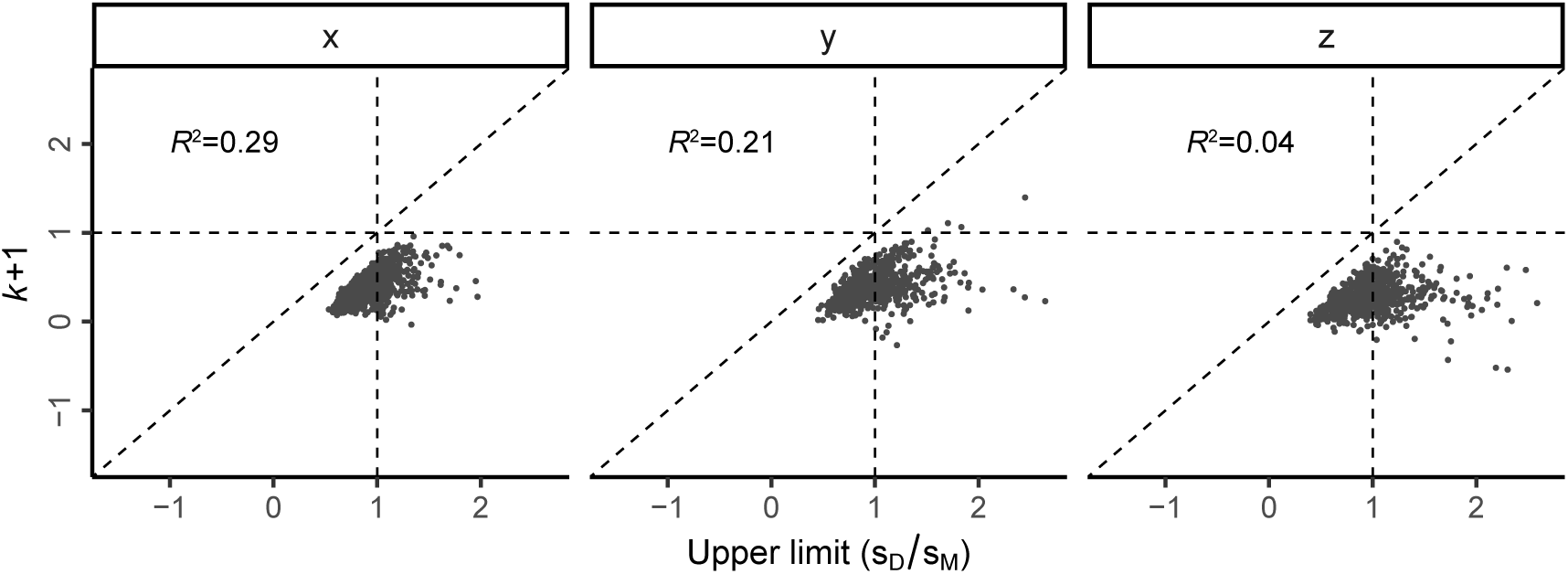
Evaluation of the σ_D_/σ_M_<1 Issue in the Estimation of *k*. In linear regression, if the independent variable has a smaller standard deviation (SD) than the dependent variable, a slope smaller than 1 will always be obtained due to the relationship between slope and correlation, *k*=σ_D_/σ_M_ × *R*. The horizontal axis shows the σ_D_/σ_M_ for each mother-daughter cell pair at three coordinates, respectively. The vertical dashed line signifies σ_D_/σ_M_=1 in each panel, indicating that the SD is overall comparable between mother and daughter cells. The vertical axis is *a*=*k*+1, and the horizontal dashed line corresponds to *k*=0 in each panel. The diagonal line is shown in each panel. Given that σ_D_/σ_M_ is the upper limit of *a*=*k*+1, all the points are distributed below the diagonal line. The small *R*^2^ in each panel suggests that *a*=*k*+1 cannot be simply explained by their upper limits.

**Fig. S11.**
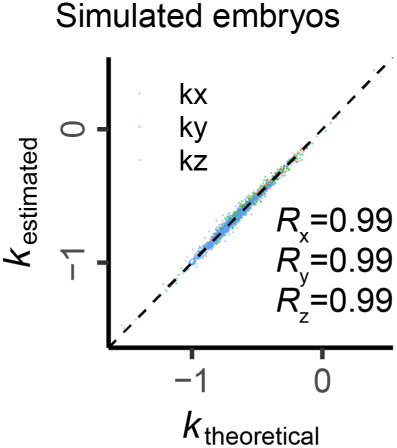
Impact of Translation-Rotation-Scaling Procedure on the Estimation of *k*. This figure compares the theoretical and observed *k* values of the three coordinates in simulated embryos. The three coordinates are labeled in different colors. The diagonal line and Pearson’s *R* for each coordinate are shown.

**Fig. S12.**
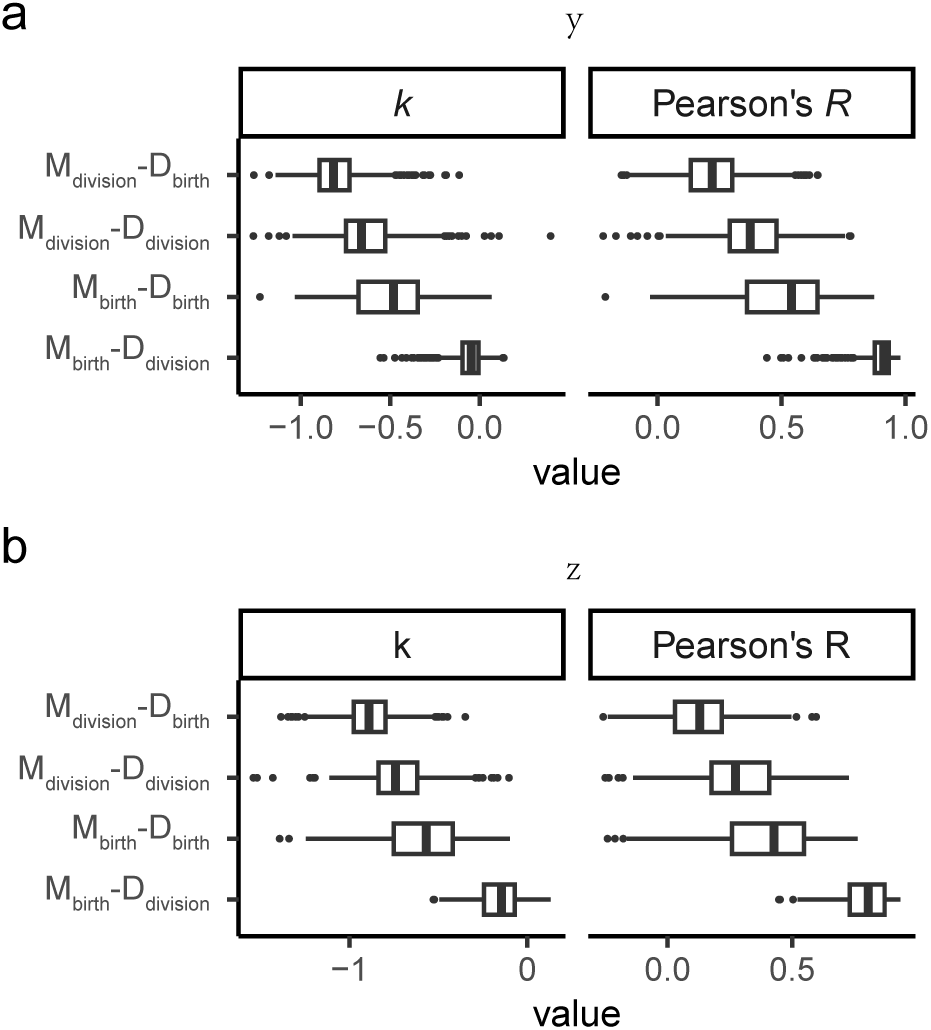
Continuous Process of Mother-Daughter Negative Feedback. This figure shows the results for the y-coordinate (a) and z-coordinate (b), corresponding to Fig. 2g.

**Fig. S13.**
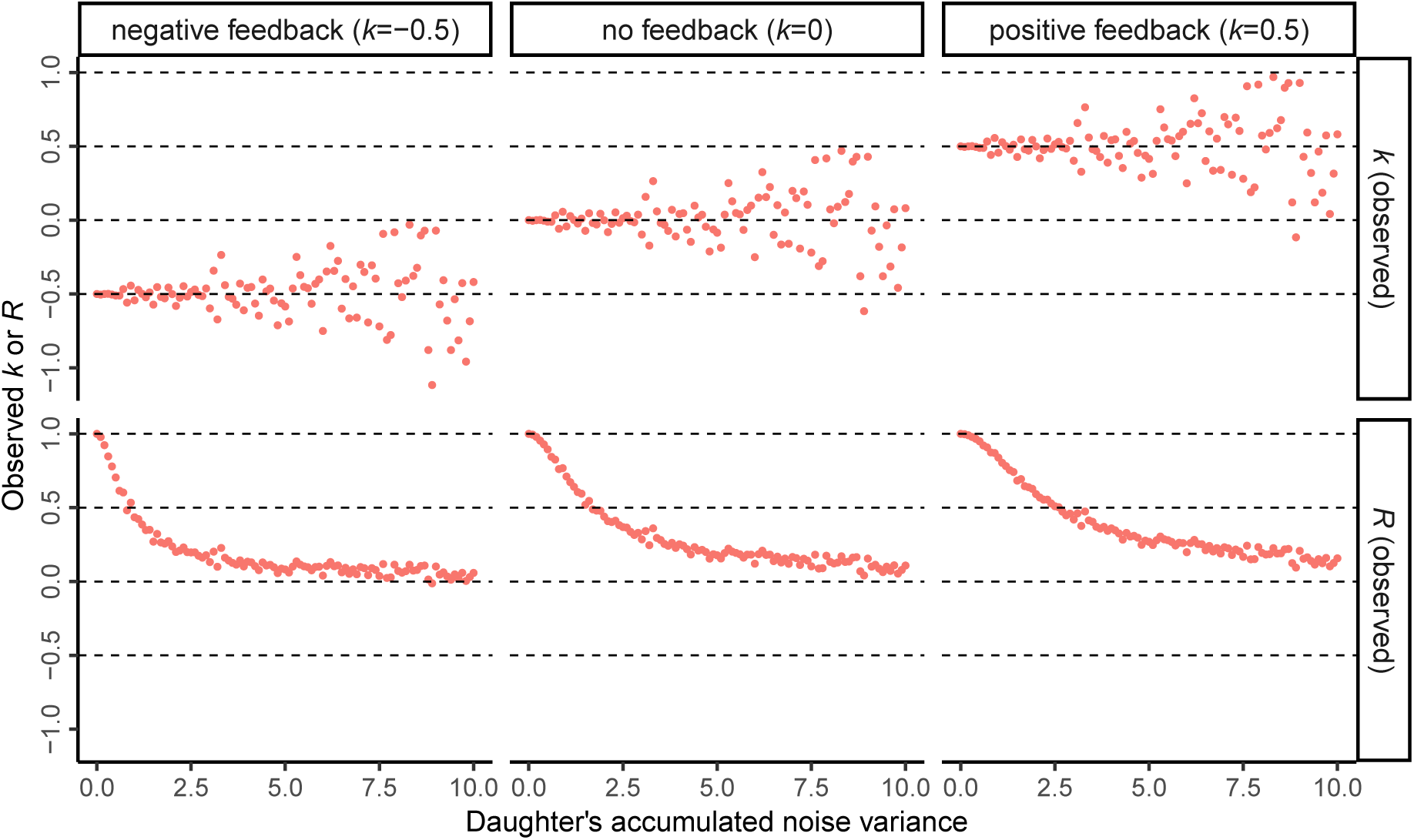
The Effect of Daughter’s Noise Accumulation on *k* and *R*. This figure illustrates how the noise accumulation in daughter cells can impact the estimated *k* values and Pearson’s *R*, based on mother’s and daughter’s noise. The process is simulated as follows: First, we generate 1,000 instances of mother’s noise from a standard normal distribution (N(0,1)). Then, the accumulated noise *ε* is generated from N(0,1) and multiplied by a factor denoting the variance level, ranging from 0 to 10. Next, the daughter’s noise is calculated using *X*_D_ = (*k*+1)*X*_M_ + *ε*, considering three different *k* values that correspond to negative feedback, no feedback, and positive feedback, respectively. For each scenario, we estimate the *k* values. The results show that the observed *R* between mother’s and daughter’s noise gradually decreases to zero as the variance of the daughter’s accumulated noise increases. However, the expected values of the observed *k* remain constant, while their variation increases. Therefore, the negative *k* values we obtained cannot simply be attributed to the accumulation of noise in the daughter cells. Considering that the daughter at birth inherits the main variance of its mother, the negative *k* values demonstrate the presence of negative feedback.

**Fig. S14.**
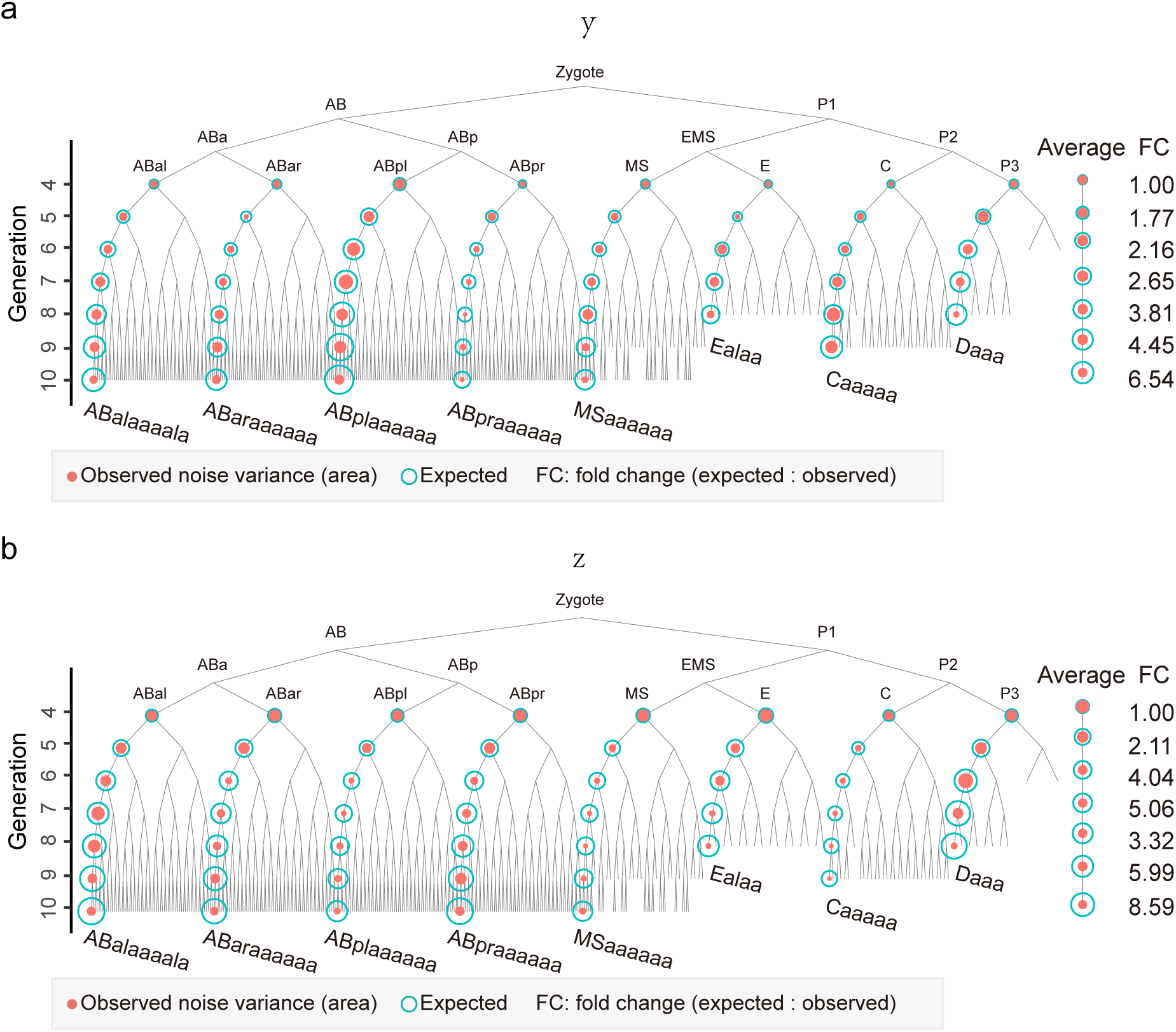
Theoretical and Observed Noise Variance Along Lineages. This figure presents the results for the y-coordinate and z-coordinate, corresponding to Fig. 2h. The theoretical and observed noise variances along different lineages are compared for these two spatial coordinates. Each lineage’s theoretical noise variance is calculated based on the noise variance of the initial cell and the estimated daughter-specific noise variance. The observed noise variance is directly measured from the data.

**Fig. S15.**
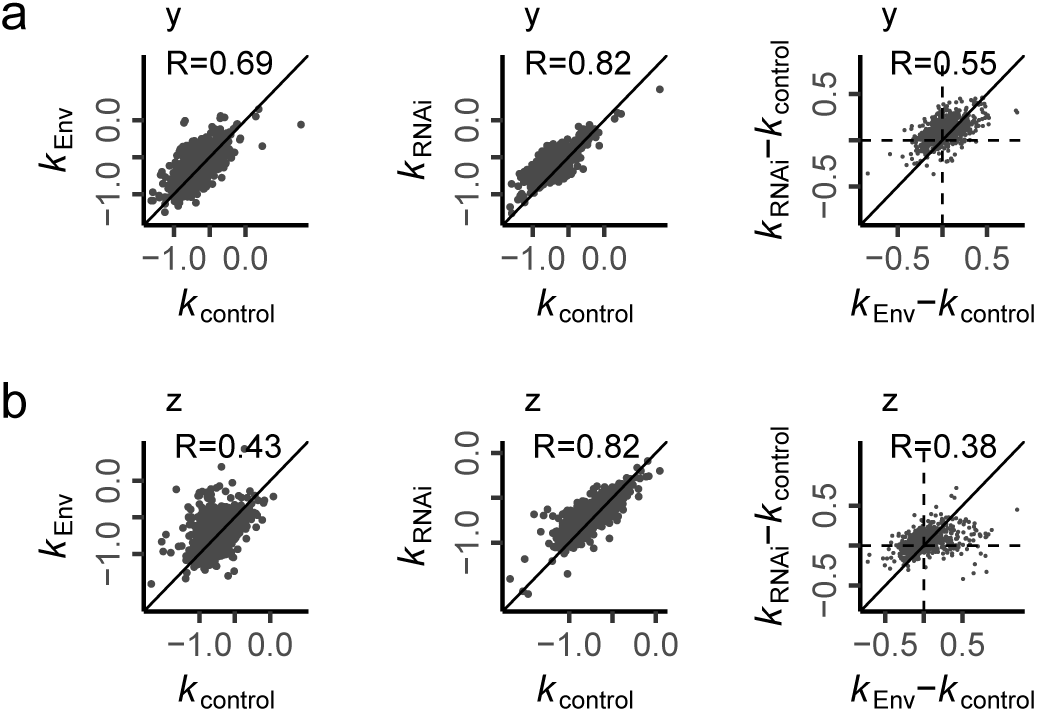
Comparison Between the *k* Values of Different Types of Embryos. These panels show the results for the y-coordinate (a) and z-coordinate (b), respectively, corresponding to Fig. 3a-c.

**Fig. S16.**
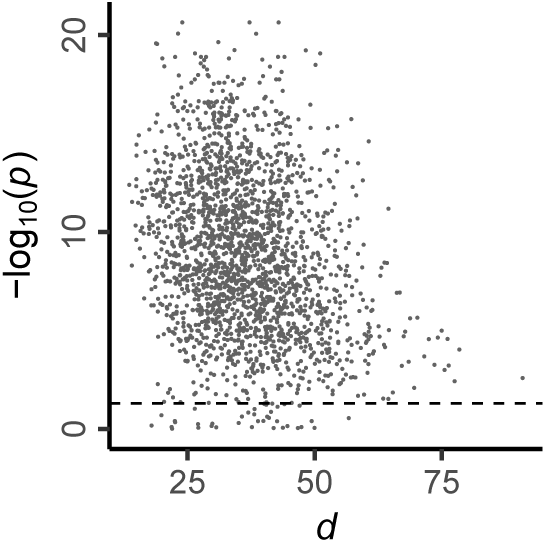
The Distribution of p-values for Noise Tolerance Level *d*. This figure demonstrates the distribution of p-values for the noise tolerance level of each cell. The noise of each cell among embryos is used to model the hatching phenotype using logistic regression. The model is formulated as logit(P)=E(*β*_0_+*β*_1_*X*), where P represents the probability of an embryo exhibiting a lethal phenotype. From this, we obtain the p-value for each coefficient using the ‘glm’ function in R. Each point in the figure represents one coordinate feature of a cell. The horizontal dashed line corresponds to a p-value of 0.05, which is commonly used as the threshold for statistical significance.

**Fig. S17.**
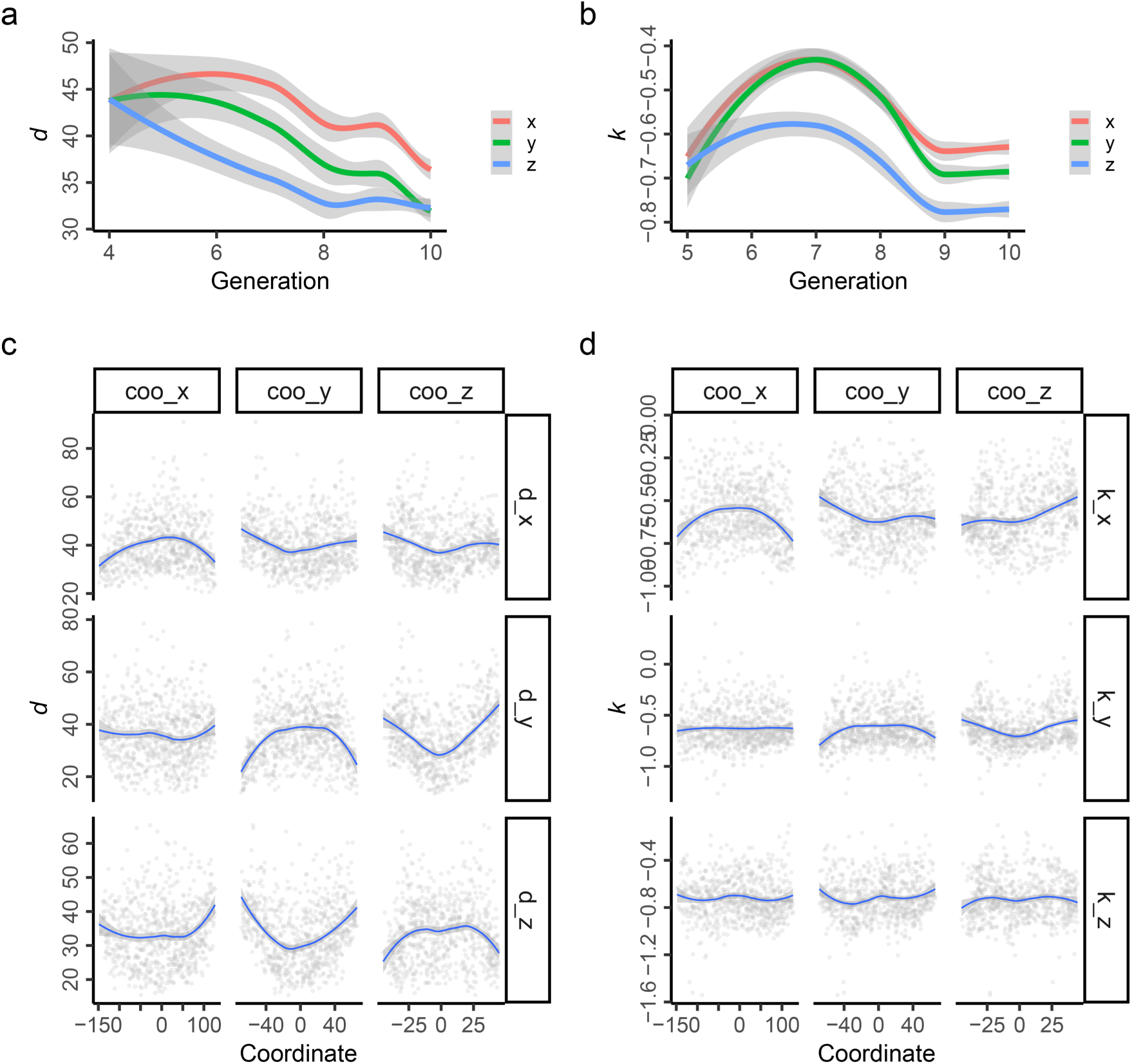
Temporal-Spatial Patterns of *k* and *d*. **(a)** This panel illustrates the decreasing trend of d values as generation increases. A loess regression is used to fit the d values against generations. The band around the fitted lines represents the 95% confidence interval. Each coordinate is represented by a different color. **(b)** This panel presents the convex patterns of *k* values across generations. **(c)** The *d* values for each coordinate are plotted against the three coordinates. Each point represents a daughter cell. The fitted line is obtained from loess regression. The three coordinates of each cell are derived from the average of 105 control embryos. **(d)** This panel is similar to (c), but considers *k* values instead.

**Fig. S18.**
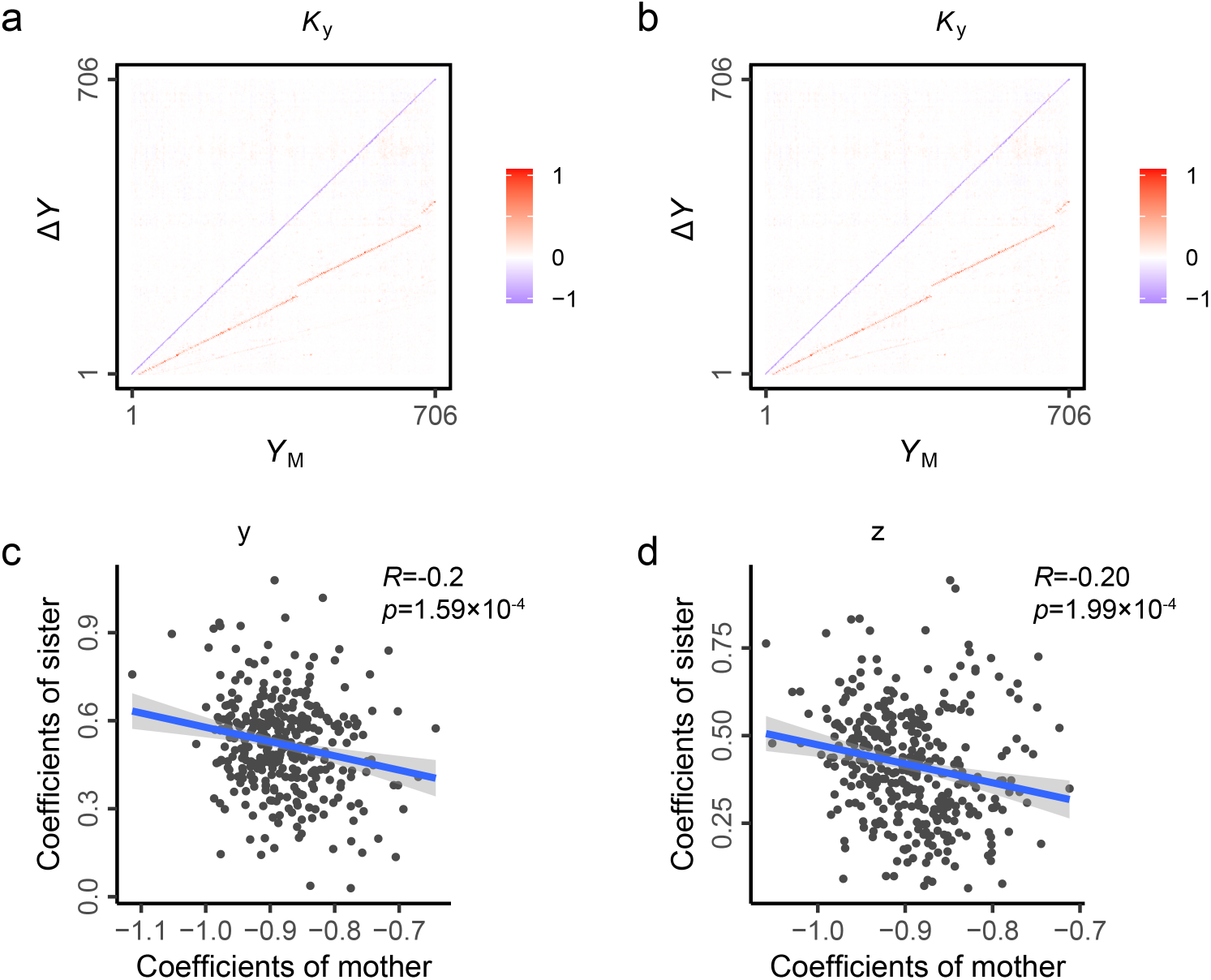
High-Dimensional Model for Position Features. This figure displays the results of high-dimensional modelling for the *y* and *z* coordinates, respectively, corresponding to Fig. 5b and Fig. 5e. **(a-b)** These heatmaps display the coefficient matrices for the high-dimensional models of the *y* and *z* coordinates, respectively. **(c-d)** These scatter plots show the negative correlation between the coefficients of mothers and sisters for focal daughters. The blue line is fitted by linear regression, with the band around the fitted lines representing the 95% confidence interval.

**Fig. S19.**
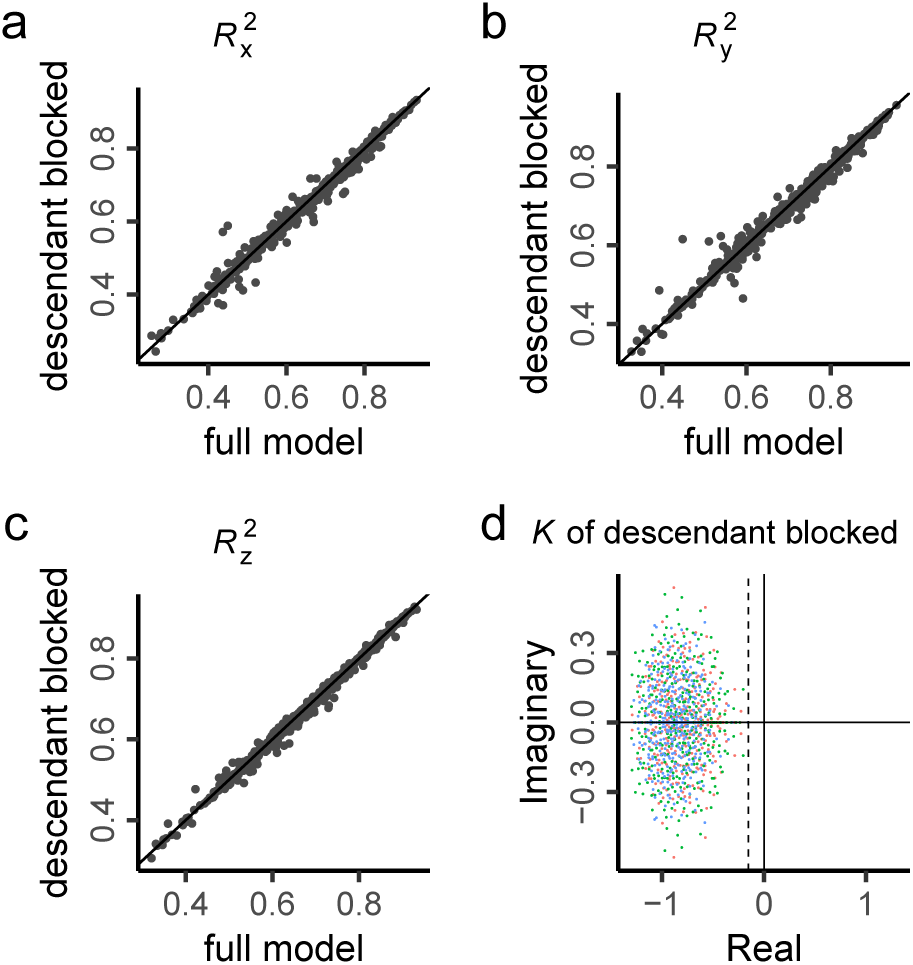
High-Dimensional Model for Position Features with Descendant Generations Blocked for Each Daughter Cell. This figure presents the high-dimensional model which includes all mother-daughter cell pairs, referred to as the full model, in comparison to the model obtained by blocking the descendant generations for each daughter cell, termed the nested model. The nested model demonstrates highly similar results to the full model. **(a-c)** These panels compare the model performance for each item in Δ*X*, Δ*Y*, or Δ*Z* between the full model and the nested model. The diagonal line is shown for reference. **(d)** This panel illustrates the distribution of eigenvalues of the nested models for *x*, *y*, and *z* coordinates, respectively. The vertical dashed line indicates the maximum real part of the eigenvalues.

**Fig. S20.**
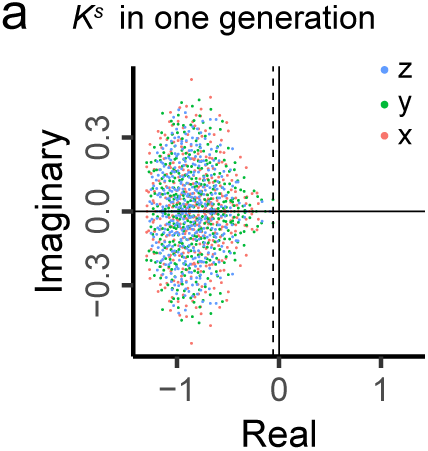
Birth-Division Stability Within a Single Cell Cycle. To assess the stability of the system from cell birth to cell division, we constructed a high-dimensional model that considers cell noise at both birth and division for each cell. The resulting coefficient matrix is termed as *K*^s^. This figure illustrates the eigenvalues of *K*^s^ for x, y, and z coordinates, respectively. The dashed line marks the maximum real part of the eigenvalues.

**Fig. S21.**
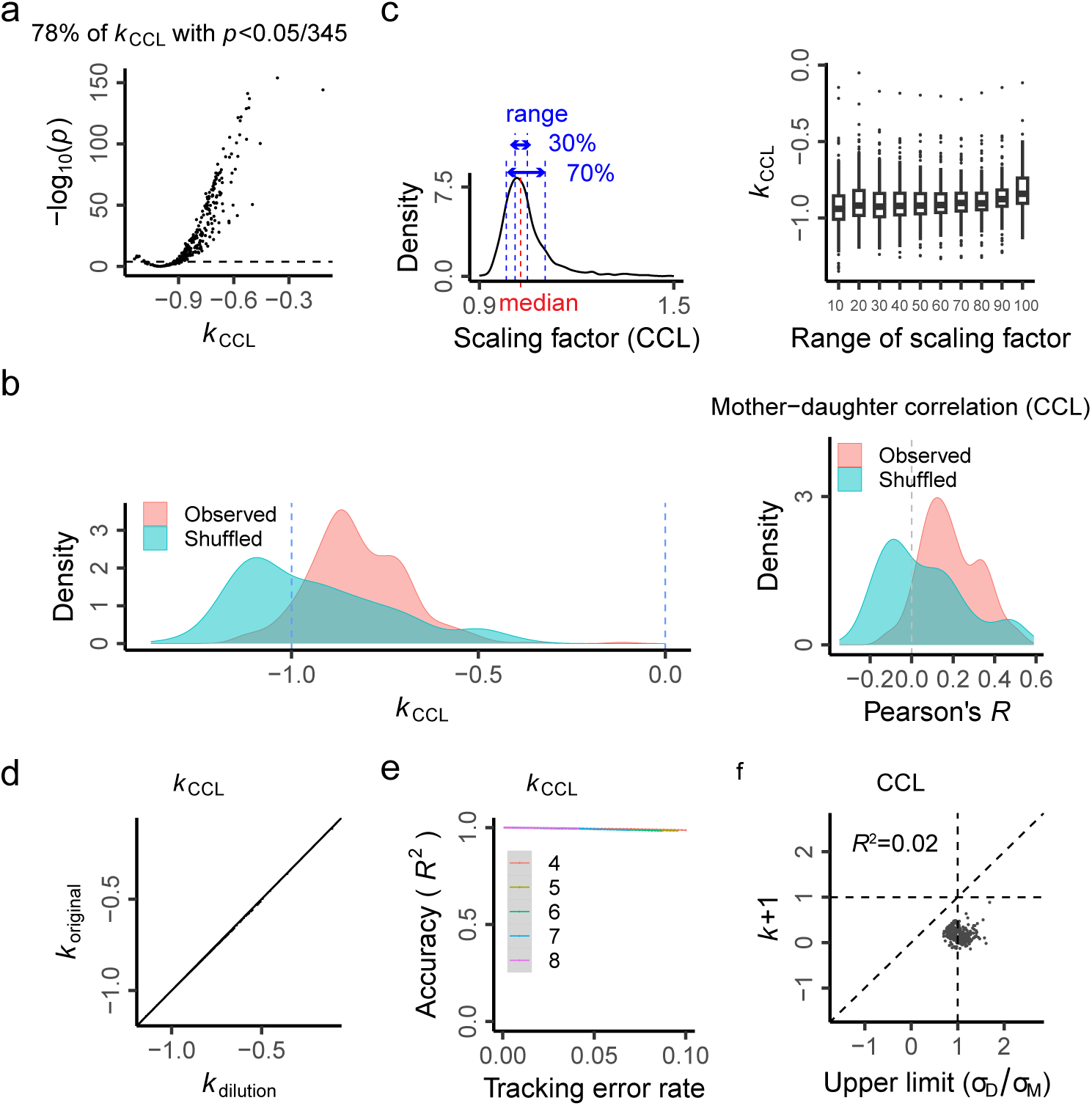
CCL’s Negative Feedback Model and Confounding Factors. **(a)** This panel shows the distribution of p-values for *k*_CCL_. **(b)** This panel compares *k* and *R* values between mother-daughter and shuffled cell pairs. **(c)** This panel displays the re-estimation of *k*_CCL_ across a series of scaling ranges, demonstrating its robustness. **(d)** This panel indicates minimal dilution effect from measurement precision on the estimation of *k* values. **(e)** This panel shows the effect of tracking error on the accuracy of *k* values with an error rate up to 10% in each generation. **(f)** This panel addresses the σ_D_/σ_M_<1 issue. In the context of CCL, this problem is equivalent to the coefficient of variation (CV) of the daughter being smaller than that of the mother before normalizing the scaled CCL noise to the average CCL of each cell. Similar to the observations in position features, the CVs of the daughter and mother cells are comparable, and their ratio only explains a minor portion of the variance in the *k* values.

**Fig. S22.**
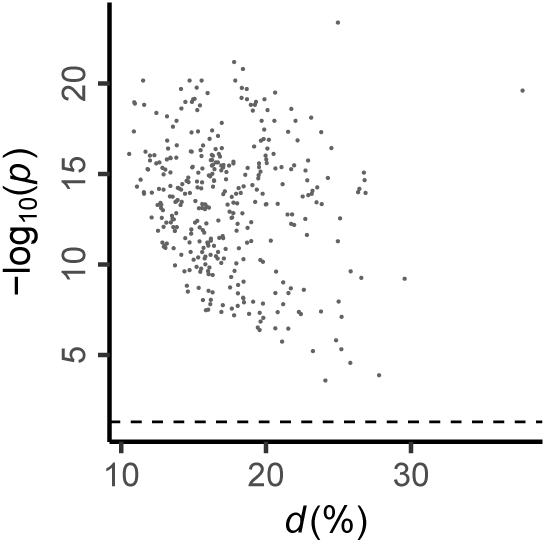
The p-values for the Noise Tolerance Level *d* of Each Cell’s CCL. This figure displays the p-values for the noise tolerance level d of each cell’s CCL. The horizontal dashed line corresponds to *p*=0.05.

**Fig. S23.**
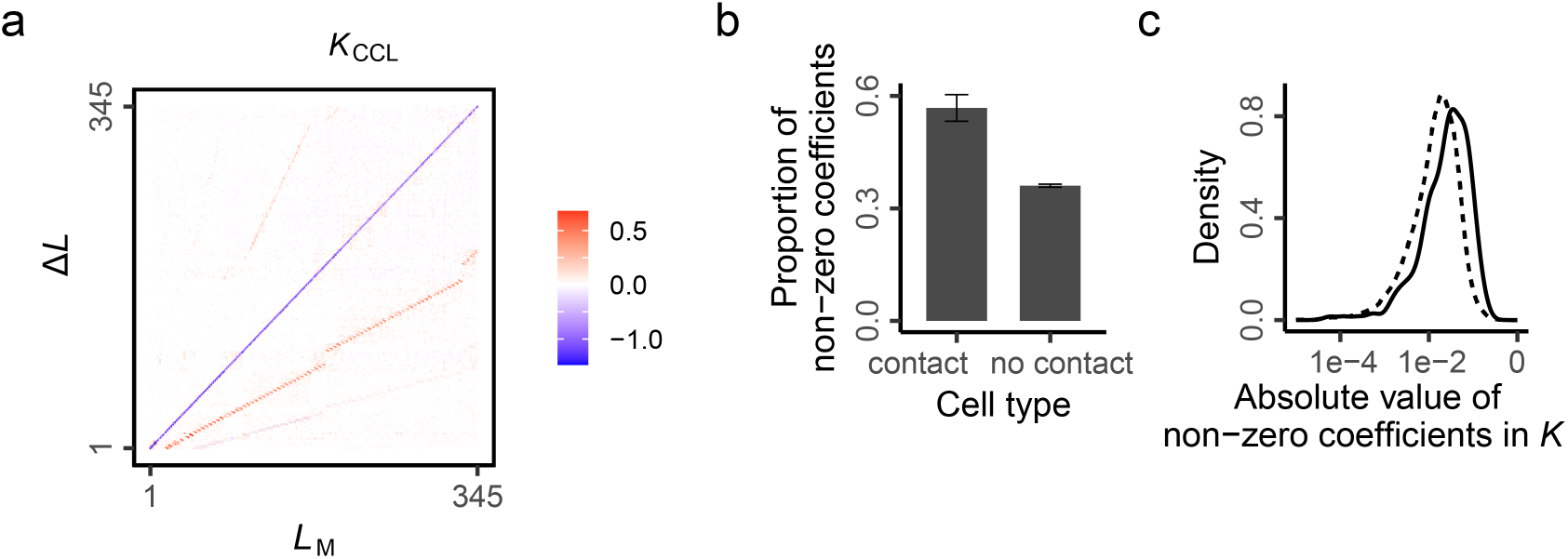
Coefficient Matrix of the High-Dimensional Model for CCL. **(a)** A heatmap displays the coefficient matrix, *K*_CCL_. **(b-c)** These panels depict the effect of cell contact in *K*_CCL_, similar to Fig. 7g-h.

**Fig. S24.**
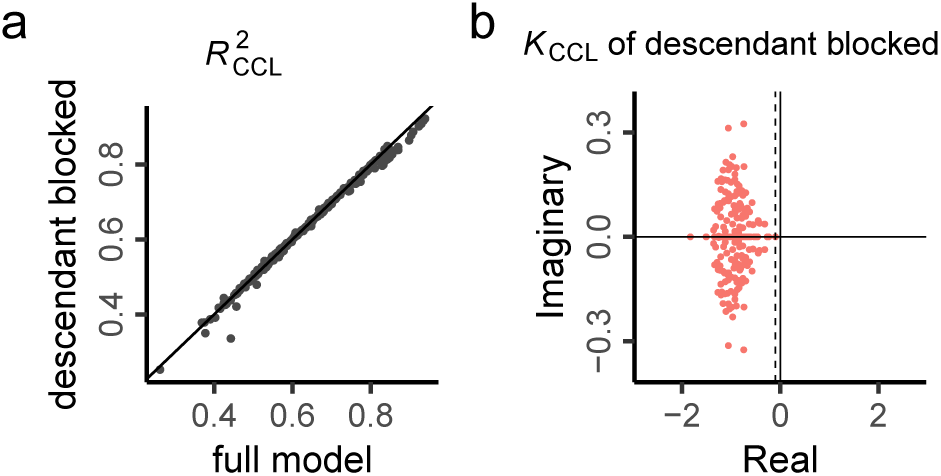
High-Dimensional Model for CCL with Descendant Generations Blocked for Each Daughter Cell. In Fig. 5a, the high-dimensional model includes all mother-daughter cell pairs (the full model). Here, we present the high-dimensional model with descendant generations blocked for each daughter cell (the nested model). The nested model shows highly similar results to the full model. **(a)** Comparison of model performance for each item in Δ*L* between the full model and the nested model. **(b)** The eigenvalues of the nested model.

**Fig. S25.**
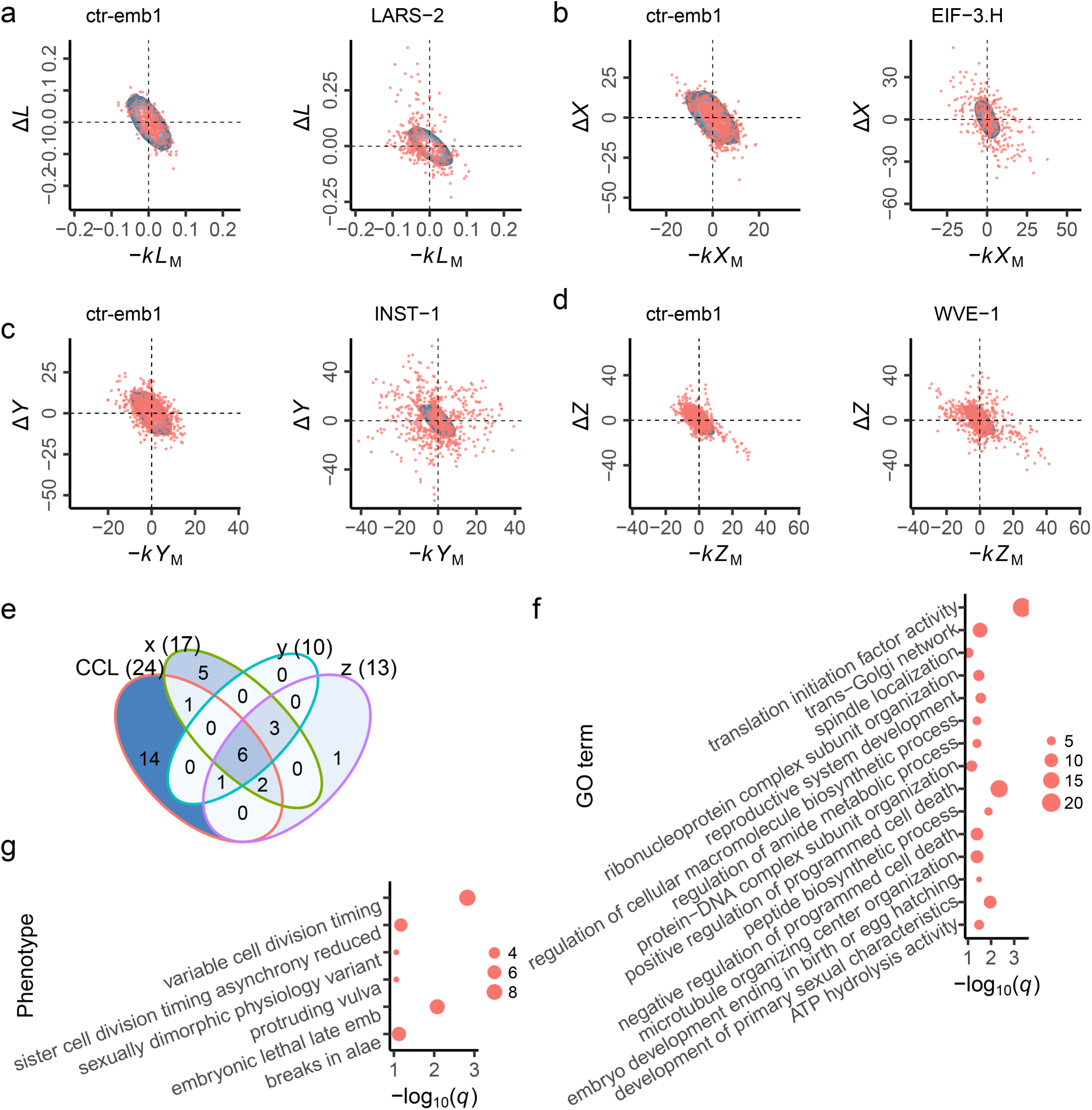
Identification of candidate canalization genes. **(a)** Illustration of CCL’s negative feedback defect induced by genetic perturbation through an RNAi experiment, comparing the distribution of (-*kL*_M_, Δ*L*) for mother-daughter (M-D) cell pairs in an example RNAi embryo (LARS−2 RNAi) with that in a control embryo (ctr-emb1). The blue background region illustrates the 90% density distribution of (-*kL*_M_, Δ*L*) for M-D cell pairs in 105 control embryos. Compared to the control embryo (ctr-emb1), the (-*kL*_M_, Δ*L*) for the example RNAi embryo deviates significantly from the expected 90% region of 105 control embryos. We expect the (-*kL*_M_, Δ*L*) for all M-D cell pairs in an embryo to align around the diagonal line across the second and the fourth quadrants—indicative of successful negative feedback (i.e., Δ*L* = *kL*_M_+*ε*)—as represented by the background blue region formed by 105 control embryos. **(b-d)** Analogous to panel (a) but for x, y, and z coordinates, respectively, depicting the distribution of (-*kX*_M_, Δ*X*) for M-D cell pairs in an example RNAi embryo (EIF-3.H RNAi), (-*kY*_M_, Δ*Y*) for M-D cell pairs in an example RNAi embryo (INST−1 RNAi), and (-*kZ*_M_, Δ*Z*) for M-D cell pairs in an example RNAi embryo (WVE−1 RNAi), in comparison to a control embryo (ctr-emb1). **(e)** A total of 33 genes are identified as candidate canalization-maintaining genes, combined together from those which display notable deviation from the mother-daughter negative feedback model (Methods). The Venn diagram shows the intersections among the candidate canalization genes identified for CCL, x, y, and z coordinates. **(f-g)** Results of GO and phenotype enrichment analyses, respectively, conducted with WormBase’s online enrichment tool. Dot size indicates fold change, and q value denotes the multiple-testing corrected p-value.

**Fig. S26.**
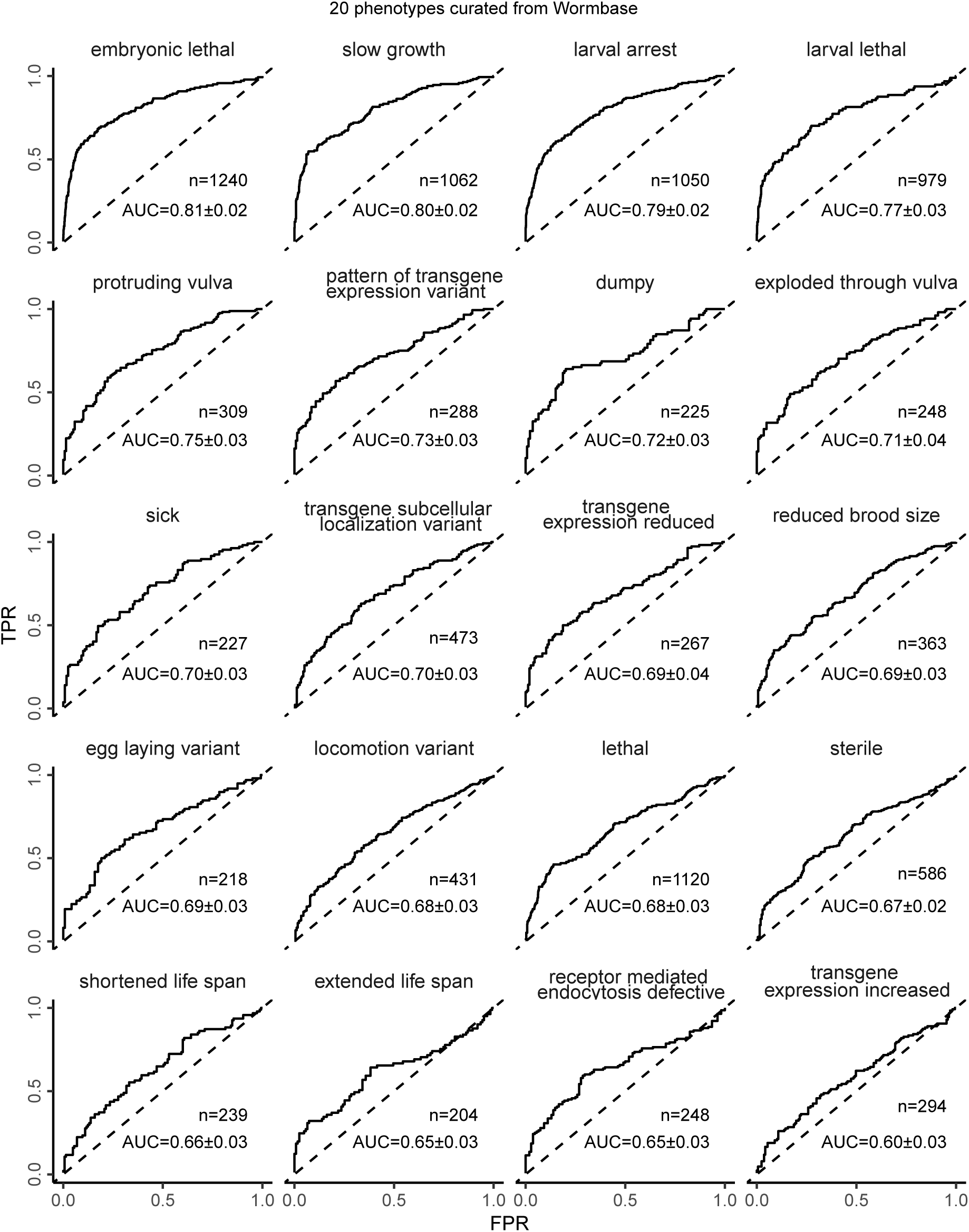
Modeling of 20 Phenotypes in WormBase by Cell Noise. We extracted the gene-phenotype annotation file from WormBase. RNAi embryos are assigned as having or lacking a phenotype based on the ‘Qualifier’ variable. An embryo is assigned a missing value when its RNAi gene does not have a record of the phenotype. We then selected the 20 phenotypes with at least 80 embryos exhibiting the corresponding phenotype for logistic regression. We used a machine learning framework similar to Fig. 4b. The Area Under the Curve (AUC) ± standard error and the number of embryos are labeled for each phenotype.

